# Pneumococcal H_2_O_2_ Reshapes Mitochondrial Function and Reprograms Host Cell Metabolism

**DOI:** 10.1101/2025.05.22.655446

**Authors:** Anna Scasny, Babek Alibayov, Ngoc Hoang, Ana G. Jop Vidal, Kenichi Takeshita, Consuelo Bautista, Eva Bengten, Antonino Baez, Wei Li, Jonathan Hosler, Kurt Warncke, Kristin Edwards, Jorge E. Vidal

**Affiliations:** Department of Cell and Molecular Biology, School of Medicine, University of Mississippi Medical Center, Jackson, MS; Department of Pharmacology and Toxicology, School of Medicine, University of Mississippi Medical Center, Jackson, MS; Center for Immunology and Microbial Research, School of Medicine, University of Mississippi Medical Center, Jackson, MS; Centro de Investigaciones en Ciencias Microbiológicas (CICM), Instituto de Ciencias (IC), Benemérita Universidad Autónoma de Puebla (BUAP), Puebla, México; Department of Physics, Emory University, Atlanta GA

## Abstract

*Streptococcus pneumoniae* (Spn), a primary cause of pneumonia, induces acute lung parenchymal damage through a unique metabolic pathway generating hydrogen peroxide (H_2_O_2_) as a byproduct. This study demonstrates that Spn-derived H_2_O_2_, primarily produced by pyruvate oxidase (SpxB), inhibits key tricarboxylic acid (TCA) cycle enzymes (aconitase, glutamate dehydrogenase, and α-ketoglutarate dehydrogenase) in lung epithelial cells, leading to citrate accumulation and diminished NADH production for oxidative phosphorylation. RNA sequencing reveals SpxB-dependent upregulation of glycolytic genes (HIF1A, IER3, HK2, PFKP), restricting pyruvate entry into the TCA cycle and increasing glucose consumption and lactate/acetate production, indicative of a Warburg-like metabolic shift that may enhance bacterial survival. Notably, mitochondrial membrane potential remains largely preserved, with minimal apoptosis despite Spn-induced stress. These findings uncover a novel mechanism of Spn-driven host metabolic reprogramming, highlighting potential therapeutic targets for pneumococcal diseases.

**Importance:** *Streptococcus pneumoniae* remains a leading cause of community-acquired pneumonia worldwide, yet the mechanisms by which it manipulates host metabolism to promote its survival and pathogenesis are not fully understood. This study reveals a novel metabolic strategy whereby pneumococcus-derived hydrogen peroxide, generated by pyruvate oxidase (SpxB), disrupts the host TCA cycle and drives a Warburg-like metabolic shift in lung epithelial cells. By inhibiting key TCA cycle enzymes and rewiring glycolytic gene expression, S. pneumoniae effectively reprograms host cell metabolism to favor its persistence, while minimizing host cell apoptosis and maintaining mitochondrial function. These insights expand our understanding of host-pathogen metabolic interactions and identify potential metabolic vulnerabilities that could be targeted to mitigate tissue damage and improve treatment outcomes in pneumococcal pneumonia.

## Introduction

*Streptococcus pneumoniae* (Spn), commonly known as the pneumococcus, is a major global health burden, particularly among children and the elderly, causing significant morbidity and mortality through pneumonia and other invasive pneumococcal diseases (IPDs)(1, 2). During pneumococcal pneumonia, lung colonization induces profound cellular damage, disrupting critical structural components such as ciliary function, β-actin architecture, and nuclear and mitochondrial DNA (3–5). Despite progress in understanding pneumococcal pathogenesis, the mechanisms driving host cell death and tissue injury remain incompletely elucidated.

A hallmark of Spn pathogenesis is its efficient metabolic activity, primarily driven by glycolysis, which supports energy and biomass production through glucose catabolism. A critical byproduct, hydrogen peroxide (H_2_O_2_), a reactive oxygen species, is increasingly implicated in host cell damage (1, 6). Spn produces H_2_O_2_ continuously via glycolysis, its primary ATP-generating pathway, with some strains generating up to 2 mM H_2_O_2_, which is detectable both extracellularly and intracellularly(7–9). Pyruvate oxidase (SpxB) and lactate oxidase (LctO) primarily drive H_2_O_2_ production, with SpxB converting pyruvate to acetyl phosphate and LctO oxidizing lactate to pyruvate, both yielding H_2_O_2_ (9, 10). SpxB accounts for ∼85% of total H_2_O_2_ production in Spn, while LctO contributes ∼15%, reflecting adaptations to oxygen-rich environments like the lungs (9, 10).

Substantial evidence implicates Spn-derived H_2_O_2_ (hereafter Spn-H_2_O_2_) in host cell death and disease severity in animal models (11–16). Our previous studies have shown that Spn-H_2_O_2_ inhibits *ex vivo* Complex I-driven respiration (CIDR)(17). Additionally, Spn infection induces mitochondrial DNA leakage in lung cells, triggering a type I interferon response (18), and in an aged murine model of pneumonia, it dysregulates mitochondrial complex expression, elevates oxidative stress, and impairs antioxidant defenses, leading to reduced ATP production (19). These findings underscore the role of H_2_O_2_ in disrupting mitochondrial function, exacerbating cytotoxicity. Furthermore, Spn alters host cell nutrient availability by depleting epithelial glycans and enhancing macrophage glutamine uptake, thereby increasing bacterial access to host-derived carbohydrates and amino acids during infection (20). These metabolic shifts fuel bacterial proliferation and inflammation, driving disease progression. Concurrently, Spn infection activates both apoptosis and pyroptosis, indicating complex cytotoxic mechanisms (3, 4, 21, 22). Our recent studies reveal that Spn-H_2_O_2_ disrupts actin and tubulin cytoskeletons in bronchial and alveolar epithelial cells, impairing structural stability and protein trafficking, leading to microtubule catastrophe and cell swelling due to loss of actin stress fibers (17, 23). This mirrors cytoskeletal damage in ventricular cardiomyocytes exposed to 100 µM H_2_O_2_, where oxidation of specific cysteine residues (Cys213, Cys295, and Cys305 in α-globin; Cys129 in β-globin) disrupts tubulin interactions, inhibiting microtubule polymerization (24–26). Moreover, Spn-H_2_O_2_ oxidizes hemoglobin *in vitro* and *in vivo*, inducing aggregation via crosslinking of α- and β-globin chains (27, 28) and generating MetOH, a product of methionine- 55 oxidation in the β-chain linked to globin collapse (23).

Mitochondria are essential for many cellular functions, with two main functions being to produce ATP through oxidative phosphorylation (OXPHOS) and regulate programmed cell death (29). The tricarboxylic acid (TCA), or Krebs, cycle, a central metabolic pathway, supplies essential intermediates such as NADH to the electron transport chain (ETC) for ATP synthesis (30). In this process, exergonic electron transfer to O_2_ is coupled to the endergonic phosphorylation of ADP to ATP via four ETC complexes (I–IV), which contain heme- or iron-sulfur (Fe-S)-containing proteins (31, 32). For instance, complexes I and III include Fe-S proteins, while complexes II, III, and IV incorporate hemoproteins with hemes b, c, and a, respectively; soluble cytochrome c (Cyt*c*) also carries heme c (31, 33). We have previously demonstrated that Spn-H_2_O_2_ oxidizes soluble Cyt*c* (*17, 34*), and this oxidant is produced intracellularly during pneumococcal invasion of lung cells (23), highlighting a direct mechanism of mitochondrial disruption.

To establish infection, Spn invades bronchial and alveolar epithelial cells, adapting to the intracellular environment for survival and proliferation. Spn’s metabolic configuration resembles that of mitochondria under stress conditions (e.g., hypoxia), with an incomplete TCA cycle lacking α-ketoglutarate dehydrogenase, which converts α-ketoglutarate to succinyl-CoA, forcing reliance on glycolysis for energy production in resource-limited niches (35, 36). Similarly, Spn possesses a partial electron transport chain (ETC), missing NADH:quinone oxidoreductase (complex I-like) and cytochrome bc1 complex (complex III-like), but encoding a minimal succinate dehydrogenase (complex II-like) for fumarate reduction, a cytochrome bd oxidase (complex IV-like) for oxygen reduction, and an ATP synthase (complex V-like) likely for proton homeostasis. This configuration, while suited for oxygen-limited environments, mirrors mitochondrial ETC reliance on cytochrome oxidases but results in continuous H_2_O_2_ production, amplifying oxidative stress in host cells. The absence of a robust TCA cycle and ETC also impairs Spn’s ability to detoxify reactive oxygen species, exacerbating host cell damage. Furthermore, Spn exploits host resources by scavenging metabolites such as amino acids and carbohydrates, upregulating glutamine uptake and depleting epithelial glycans to fuel its metabolism and enhance virulence (20). These metabolic adaptations highlight Spn’s ability to disrupt host cellular processes through strategic reprogramming, motivating this study to investigate the H_2_O_2_-mediated effects on host mitochondrial function and metabolism during pneumococcal infection.

## RESULTS

### Oxidation of Cyt*c* by *S. pneumoniae-*H_2_O_2_ creates a tyrosyl radical

Spn-H_2_O_2_ is known to oxidize heme groups in hemoglobin and Cyt*c,* key components of the mitochondrial ETC (17, 23, 34). To investigate the kinetics of H_2_O_2_-mediated Cyt*c* oxidation, Todd-Hewitt broth supplemented with yeast extract (THY) was enriched with 56 µM Cyt*c* (THY-Cyt*c*) and inoculated with Spn. Supernatants from time-course experiments were analyzed by spectroscopic methods to assess oxidation dynamics (37, 38). Compared to mock-inoculated THY-Cyt*c* cultures, statistically significant heme degradation was observed in those incubated with TIGR4 for 6 h (Fig. 1A) and in those inoculated with EF3030 and incubated for 4 and 6 h (Fig. 1C). In mock-inoculated THY-Cyt*c* cultures incubated for up to 6 h, the Soret peak of heme-Fe^3+^ exhibited a maximum absorbance at 405 nm (Fig. 1B and 1D). In contrast, heme degradation observed in cultures of Spn strains, TIGR4 (Fig. 1B) or EF3030 (Fig. 1D) was characterized by the significant loss of the Soret peak and the appearance of a different species absorbing at 325 nm (Fig. 1B and 1D, arrow). Heme degradation and the formation of the 305 nm species were not detected in cultures of wt strains treated with catalase or in cultures of strains deficient in Spn-H_2_O_2_ production such as TIGR4Δ*spxB*Δ*lctO* or EF3030Δ*spxB*Δ*lctO* (Fig. 1B and 1D).

**Figure 1.**
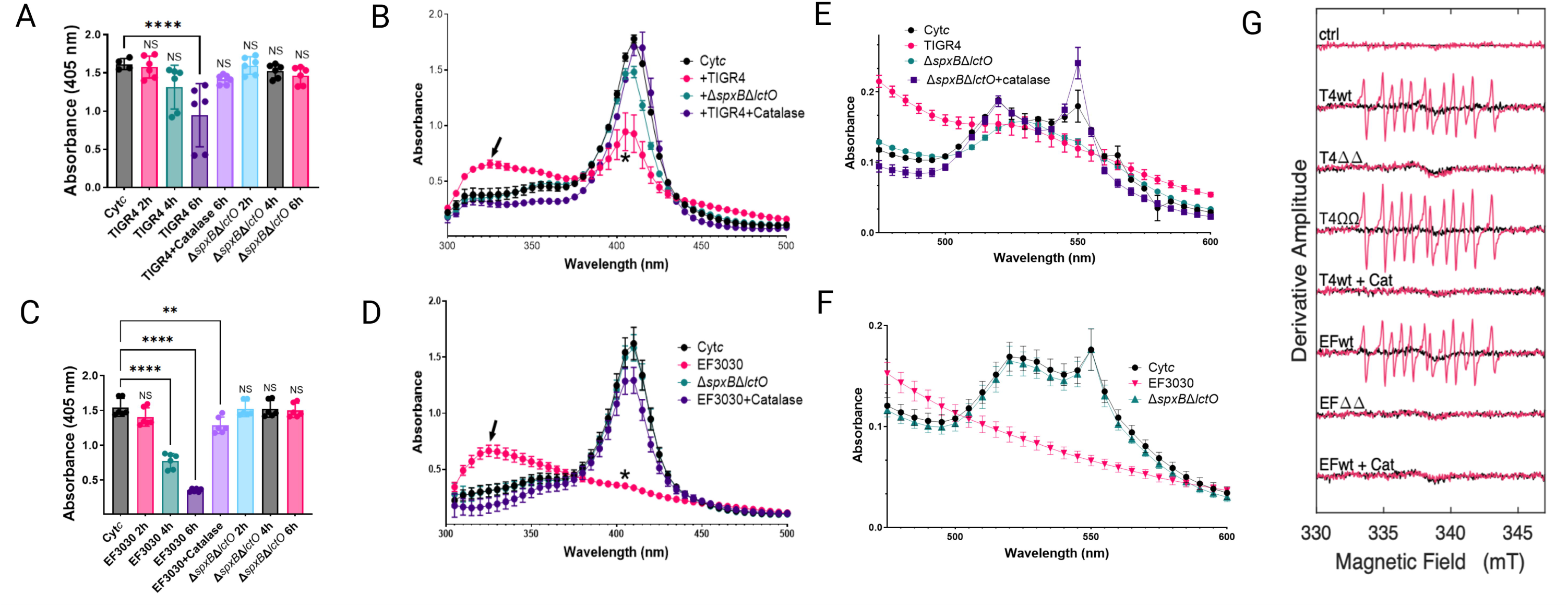
Spn-H_2_O_2_ induces cytochrome c oxidation and tyrosyl radical formation. (A-F) Spn strains were incubated in THY-Cyt*c* medium at 37°C with 5% CO_2_. At the indicated time (A, C) or 6 h post-inoculation (B, D, E, F), supernatants were analyzed by spectroscopy. Controls included mock-infected medium (Cyt*c*), or TIGR4 and EF3030 supplemented with 500 U catalase. Panels (A, C) show 405 nm absorbance, (B, D) show Soret peak absorbance, and panels (E, F) display α and β peak absorbance. Data represent means ± standard error (SE) from N=3 independent experiments, each with two technical duplicates. Statistical significance was assessed using one-way ANOVA with Dunnett’s post hoc test (A, C) or Student’s t-test (B, D). Significance levels: NS, not significant; *p < 0.05; **p < 0.001; ***p < 0.0005; ****p < 0.0001. (G) For spin-trapping, supernatants from Spn strains cultured for 4 h in THY medium were used. Cytochrome c (Cytc, 56 μM) and DEPMPO (30 mM) were added to supernatants to a final volume of 40 μL. Spectra (-Cyt*c*, black; +Cyt*c*, red) represent an average of 20 scans, corrected for buffer baseline. Uninfected THY medium served as a control (ctrl). Spn-H_2_O_2_-free samples (+Cat) were prepared by pretreating supernatants with catalase for 1 min.

The α and β absorption peaks of Cyt*c*, observed at 550 and 520 nm, respectively (39, 40) underwent oxidation in 6 h cultures of TIGR4 (Fig. 1E) and EF3030 (Fig. 1F) indicative of porphyrin ring degradation. Notably, TIGR4Δ*spxB*Δ*lctO* retained residual oxidative activity on the α and β peaks, possibly via minor SpxB/LctO-independent pathways. This residual activity was completely abrogated upon catalase treatment (Fig. 1E), underscoring the role of H_2_O_2_ in driving heme degradation in these strains.

Previous studies have shown that H_2_O_2_ oxidation of hemoproteins generates highly reactive and toxic radicals, including the ferryl radical (^•^Hb^4+^), and tyrosyl radical (5, 41–44). To determine whether radical species form during Cyt*c* oxidation by Spn-H_2_O_2_, pneumococcal strains TIGR4, EF3030, their Δ*spxB*Δ*lctO* mutants, and a complemented strain (Ω*spxB*-*lctO*) were cultured in THY medium. Supernatants were combined with Cyt*c* and the spin trap DEPMPO [5-(Diethoxyphosphoryl)-5-methyl-1-pyrroline-N-oxide, 98%], and radical formation was monitored using continuous wave electron paramagnetic resonance (EPR). EPR spectra from TIGR4, TIGR4Ω*spxB*-*lctO*, and EF3030 supernatants with Cyt*c* were identical (Fig. 1G), indicating tyrosyl radical formation from Cyt*c* oxidation. No EPR signal was detected in Δ*spxB*Δ*lctO* mutant supernatants or wild-type cultures treated with catalase. The observed EPR spectra are consistent with both protein-attached and free-solution carbon-centered radicals. Spn strains TIGR4 and EF3030 induce significantly elevated intracellular ROS levels in lung epithelial cells compared to their isogenic Δ*spxB*Δ*lctO* mutants or mock-infected controls (45, 46), supporting a link between extracellular tyrosyl radical formation and intracellular oxidative stress.

### *S. pneumoniae-*derived H_2_O_2_ contributes to mitochondrial aggregation and apical migration within alveolar cells

Disruption of soluble Cyt*c* increases reactive oxygen species (ROS) production and triggers apoptotic cell death, which can induce mitochondrial morphological changes through fusion or fission events, altering organelle structure by combining or dividing mitochondria, respectively (47). To investigate the impact of Spn infection on mitochondrial structure, human alveolar A549 cells were infected with pneumococcal strains TIGR4, EF3030, and their Δ*spxB*Δ*lctO* mutants for 8 h. Cells were stained for mitochondria (green), DNA (blue), and actin (red), and analyzed by confocal microscopy (Fig. 2).

**Figure 2.**
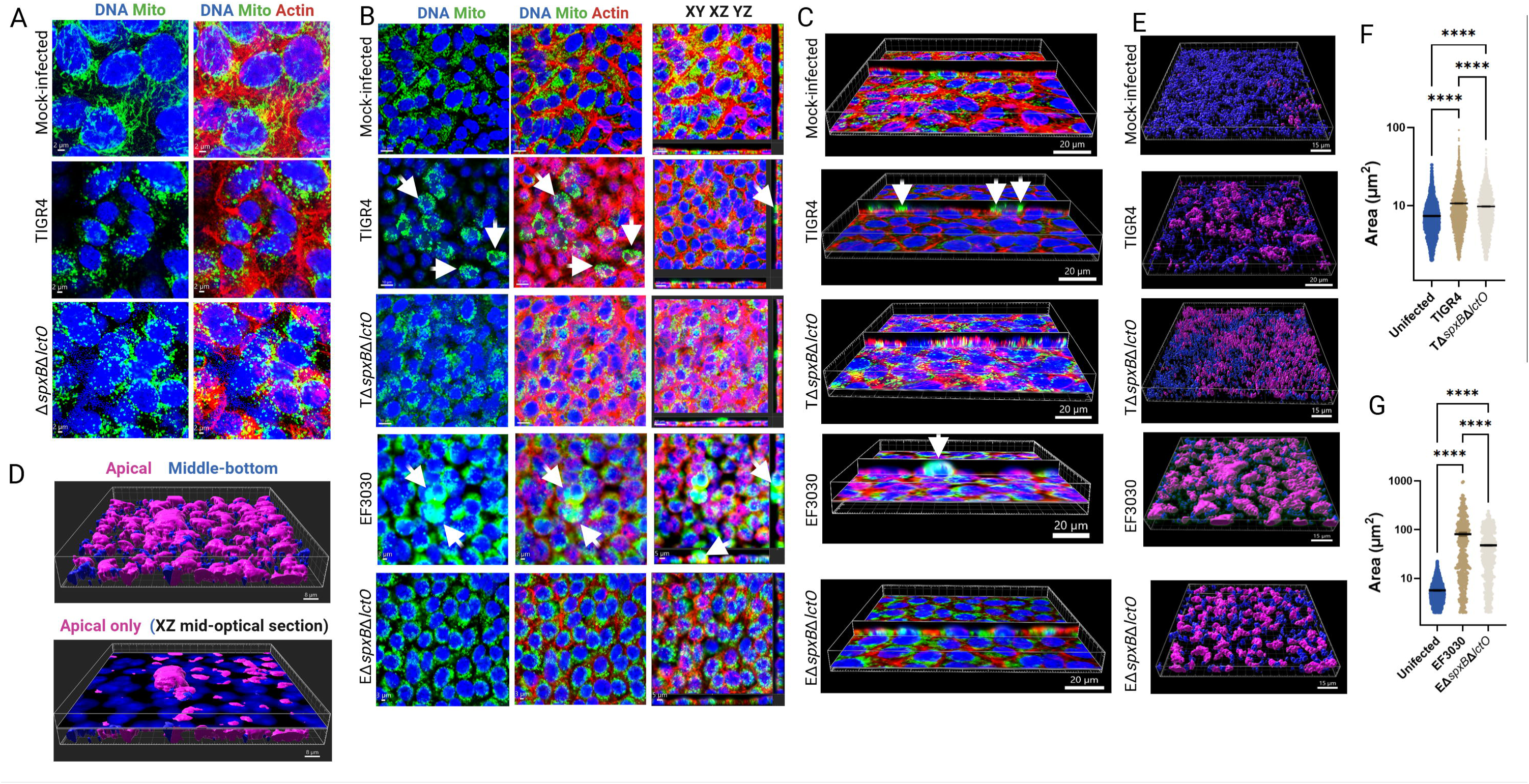
Spn-H_2_O_2_ modifies mitochondrial morphology, induces aggregation, and promotes mislocalization. Human alveolar A549 cells were either mock-infected or infected with Spn strains and incubated for 8 h at 37°C with 5% CO_2_. Infected cells were stained for mitochondria (green), DNA (blue), and actin (red), imaged via confocal microscopy, and analyzed using Imaris software. Panels (A and B) display XY optical-middle sections (0.5 µm) from z-stacks. Panel (B) additionally includes XY, XZ (bottom), and YZ (side) optical sections. Three-dimensional reconstructions were generated from z-stacks, with panel (C) presenting simultaneous XY and XZ optical sections. Arrows in panels (B, C) indicate aggregated mitochondria. Confocal microscopy images are representative of at least three independent experiments. (D-G) Imaris software was employed to generate masks from the mitochondrial channel, enabling the assessment of subcellular localization (apical, middle, or basal) and quantification of mitochondrial area (µm²). Panel (D) depicts cells infected with strain EF3030, with the top panel showing all z-stacks, highlighting predominant apical mislocalization of mitochondria, and the bottom panel a middle optical section. Panel (E) presents three- dimensional images across all conditions as in (D), while panel (F and G) quantifies mitochondrial area under each infection condition. Data are expressed as mean ± SEM from confocal micrographs of three independent experiments. Statistical significance was assessed using one-way ANOVA with Dunnett’s post hoc test, and **** indicating *p*< 0.0001.

In mock-infected control A549 cells, mitochondria exhibited a well-developed, interconnected network of elongated, filamentous structures surrounding the nuclei, characteristic of healthy, actively respiring cells (Fig. 2A). In contrast, both TIGR4-, and TIGR4Δ*spxB*Δ*lctO*-infected alveolar cells displayed fragmented and punctate mitochondria with a diffuse cytoplasmic distribution. These mitochondria were significantly smaller than those in mock-infected cells and exhibited irregular morphologies (Fig. 2A).

At different magnifications, A549 cells infected with Spn wild-type strains TIGR4 or EF3030 exhibited rounding and aggregation, with clusters of mitochondria forming around the nuclei (Fig. 2B, arrows). A more pronounced aggregation of mitochondria was apparent in EF3030-infected cells (Fig. 2B, arrows). Three-dimensional imaging analysis revealed that these mitochondrial clusters were predominantly localized on the apical surface of the infected cells (Fig. 2B and 2C). In contrast, mitochondria in TIGR4Δ*spxB*Δ*lctO*- and EF3030Δ*spxB*Δ*lctO*-infected alveolar cells appeared unevenly distributed, with varying sizes and intensities, suggesting aggregation around certain areas, near the nuclei or in regions of cellular stress (Fig. 2B and 2C). Similar mitochondrial aggregation was observed in human bronchial Calu-3 cells infected with Spn strains (Fig. S1).

Employing three-dimensional analysis, we categorized mitochondria in mock-infected and Spn-infected cells based on their spatial proximity to the apical surface. Mitochondria located within 5 µm of the cell surface were classified as ’apical,’ while those positioned deeper within the cellular architecture were designated as ’middle-bottom’ (Fig. 2D). Quantitative assessment revealed that the area of mitochondrial aggregates at the apical side of TIGR4- and EF3030-infected alveolar epithelial cells (Fig. 2E) was significantly greater than that observed in mock-infected cells or cells infected with hydrogen peroxide-deficient knockout mutants (Fig. 2F and 2G) highlighting the impact of Spn-H_2_O_2_ on mitochondrial clustering and distribution. Taken together, these results indicate that Spn-derived H_2_O_2_ induces mitochondrial aggregation and apical migration in infected alveolar cells, likely exacerbating oxidative stress by disrupting mitochondrial dynamics.

### *S. pneumoniae* inhibits mitochondrial respiration in host cells

To investigate the impact of Spn-H_2_O_2_ on mitochondrial respiration, we employed an *ex vivo* model of infection. Rat heart mitochondria were isolated and subsequently incubated with supernatants from TIGR4, EF3030, or its H_2_O_2_-deficient mutant, Δ*spxB*Δ*lctO*, following 4 hours of growth in THY. Hydrogen peroxide levels in the supernatants were then quantified. Strain TIGR4 produced a median H_2_O_2_ concentration of 499.5 µM, while EF3030 secreted 1,119 µM H_2_O_2_ into the supernatant (Fig. 3A). The H_2_O_2_ levels in the supernatants of the mutant derivatives were significantly lower (<8.86 µM) (Fig. 3A).

**Figure 3.**
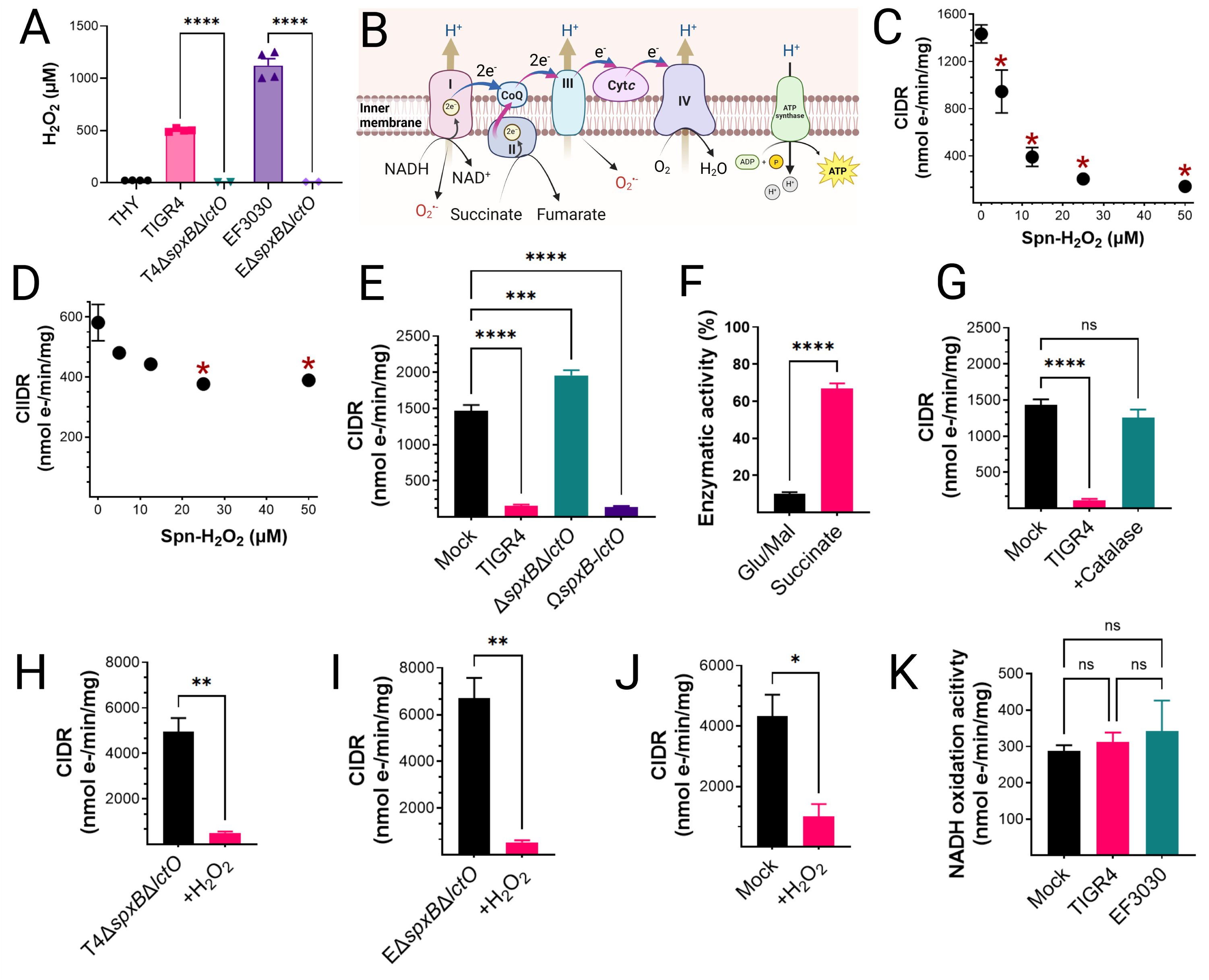
Selective inhibition of complex I-driven respiration by Spn-H_2_O_2_. (A) H_2_O_2_ concentrations in supernatants from Spn cultured in THY medium (37°C, 4 h) were quantified using Amplex Red assay (LOD<0.56 µM). Data represent mean ± SEM; unpaired Student’s t- test, ****p < 0.0001. (B) Schematic of the mitochondrial ETC in lung epithelial cells(68). (C-E) Heart mitochondria were treated with filter-sterilized Spn culture supernatants from Spn strains collected after 4 h of growth. (C, E, G-J) Complex I-driven respiration (CIDR) and (D) Complex II-driven respiration (CIIDR) were measured via Oroboros O2K FluoRespirometry. (F) Residual Complex I and II enzymatic activities relative to mock-treated controls. (G) CIDR in mitochondria treated with TIGR4 supernatants with or without catalase. (H-I) CIDR in mitochondria treated with TIGR4 Δ*spxB*Δ*lctO*, or EF3030 Δ*spxB*Δ*lctO* supernatants with or without exogenous H_2_O_2_ (50 µM), compared to mock-treated controls. (J) CIDR inhibition by exogenous H_2_O_2_ (50 µM). (K) Complex I activity in mitochondria treated with Spn supernatants (∼50 µM H_2_O_2_) or mock-treated controls, measured as NADH oxidation rate normalized to protein content. Data represent mean ± SEM; (F, H-J) unpaired Student’s t-test, *p < 0.05; (C-E, G, K) one-way ANOVA with Dunnett’s post hoc test, ns, not significant; *p < 0.05; **p < 0.005; ****p < 0.0001.

Oxygen consumption, a proxy for electron transference through CIDR, or complex-II driven respiration (CIIDR) (Fig. 3B), was investigated using a high-resolution Oroboros O2k FluoRespirometer. Isolated mitochondria were treated with the TCA cycle substrates glutamate and malate to generate NADH and initiate CIDR, which facilitates the transfer of electrons through complexes I, III, and IV (Fig. 3B). Alternatively, mitochondria were treated with succinate to drive electron transfer through CIIDR (29, 33) (Fig. 3B). Notably, mitochondrial CIDR was significantly inhibited by supernatants from TIGR4 at 5 µM H_2_O_2_, with maximal inhibition of mitochondrial respiration observed at concentrations ≥12.5 µM (Fig. 3C). In contrast, CIIDR was not inhibited by TIGR4 cultures containing the lowest H_2_O_2_ concentration, but statistically significant inhibition occurred at concentrations ≥25 µM (Fig. 3D). The factor in the supernatant causing the inhibition of mitochondrial respiration was Spn-H_2_O_2_, as an equivalent volume of supernatant from the TIGR4Δ*spxB*Δ*lctO* mutant did not inhibit CIDR, whereas the supernatant from the complemented strain caused CIDR inhibition (Fig. 3E). A comparison of residual CIDR and CIIDR activity revealed a marked difference between mock-infected and TIGR4-treated mitochondria. Mitochondria exposed to TIGR4 cultures exhibited ∼10% of the CIDR activity observed in the mock-infected control, while CIIDR activity remained significantly higher (Fig. 3F). These findings suggest that Spn-H_2_O_2_ targets complex I, the NADH dehydrogenase complex, or TCA cycle enzymes involved in NADH synthesis.

To determine whether Spn-H_2_O_2_-mediated inhibition of CIDR is reversible or irreversible, we performed a catalase rescue assay and an *in vitro* reduction treatment. Incubating TIGR4-treated mitochondria with catalase fully restored CIDRy (Fig. 3G), whereas treating mitochondria exposed to Δ*spxB*Δ*lctO* supernatant with exogenous H_2_O_2_ significantly reduced CIDR (Fig. 3H, 3I). Furthermore, exogenous H_2_O_2_ treatment inhibited CIDR in these mitochondria (Fig. 3J). These results indicate that Spn-H_2_O_2_-induced inhibition of CIDR reversible and redox-sensitive, rather than resulting from irreversible oxidative damage.

### *Streptococcus pneumoniae*-mediated inhibition of mitochondrial respiration occurs independently of electron transport chain activity

To further investigate the impact of Spn on mitochondrial function, we assessed the oxidative activity of individual complexes within the ETC. This was achieved by measuring the activity of each complex in mitochondria exposed to either sterile bacterial culture medium (mock) or supernatants from Spn strains TIGR4 and EF3030. Notably, no significant differences were observed in the oxidative capacity of complex I for its substrate NADH (Fig. 3K), or in the activity of any other ETC complexes, between mitochondria exposed to bacterial culture medium and those exposed to Spn supernatants (Fig. S2A-C). Therefore, inhibition of mitochondrial respiration by Spn occurs upstream the ETC.

### *S. pneumoniae* H_2_O_2_-mediated inhibition of TCA cycle enzymes downstream of malate dehydrogenase in mitochondrial respiration

Our results demonstrate that NADH, the substrate for complex I, can be oxidized sustaining mitochondrial respiration in the presence of Spn cultures (Fig. 3K). However, when mitochondria are supplemented with TCA cycle substrates that generate NADH, Spn cultures inhibit respiration. These findings led us to hypothesize that Spn-H_2_O_2_ selectively oxidizes TCA cycle enzymes, thereby impairing NADH production. To assess this, we stimulated cellular oxygen consumption in CIDR with a panel of TCA cycle substrate mixtures: pyruvate and malate, glutamate and malate, or succinate alone (Fig. 4A). It is well established that the substrate combinations optimize electron flow into complex I enhancing OXPHOS capacity (48). Mitochondrial respiration driven by substrates pyruvate and malate was not significantly inhibited by cultures of Spn (Fig. 4B). Similarly, cellular oxygen consumption stimulated by incubation with succinate (via complex II) was unaffected by cultures of the pneumococcus. Conversely, Spn strains TIGR4 and EF3030 significantly inhibited mitochondrial respiration induced by glutamate and malate (Fig. 4B). This data supports the hypothesis that Spn cultures inhibit enzymes of the TCA cycle, specifically downstream of citrate synthase (CS), which catalyzes the irreversible conversion of acetyl-CoA and oxaloacetate to citrate leading to citrate accumulation in TIGR4-infected Calu-3 cells, consistent with findings by Surabhi et al. (2021)(49).

**Figure 4.**
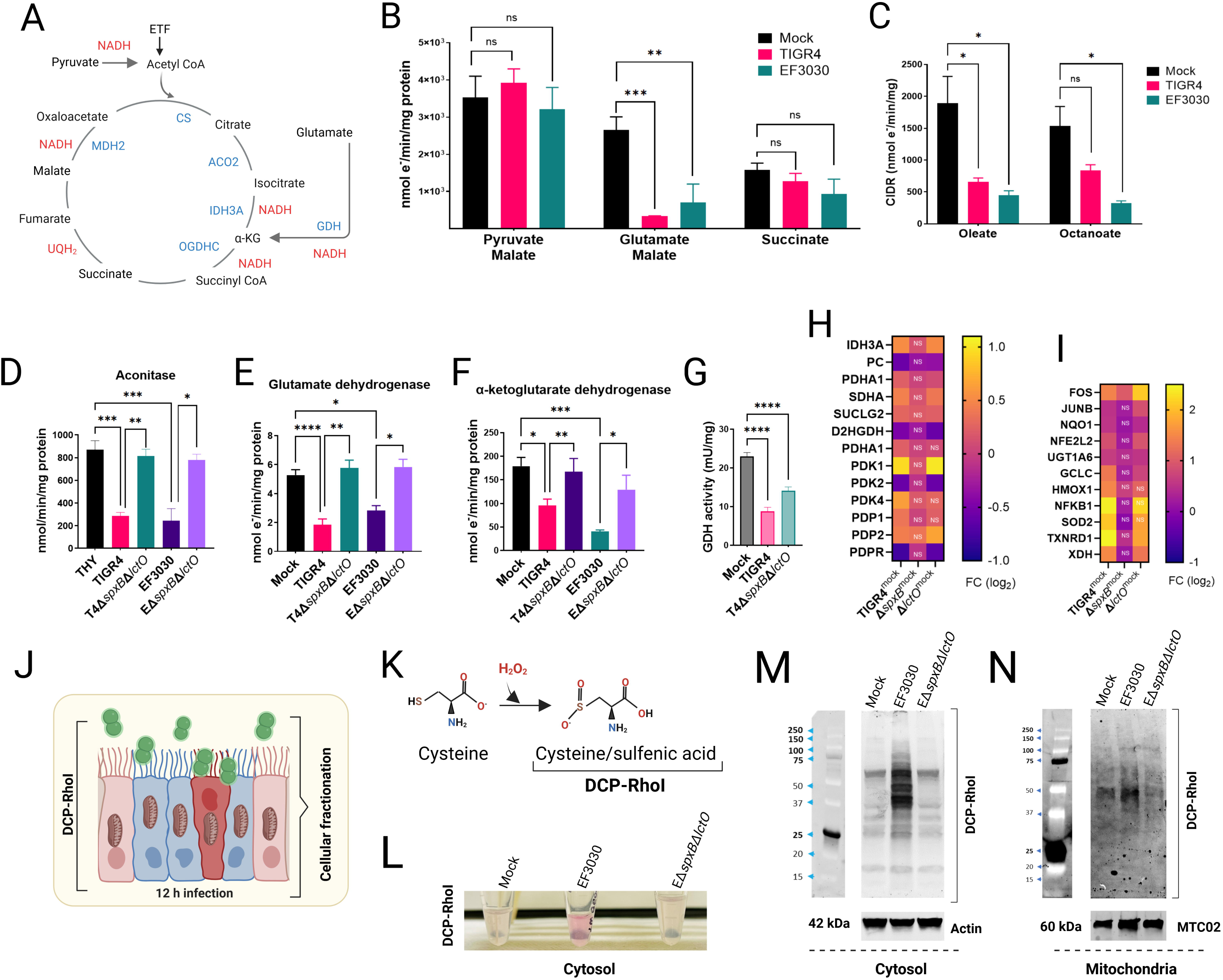
*Streptococcus pneumoniae*-derived. **H**_2_**O**_2_ **inhibits TCA cycle enzymes and mitochondrial respiration.** (A) Diagram of the canonical tricarboxylic acid (TCA) cycle, highlighting enzymes and electron donors (NADH, FADH_2_). ETF, electron-transferring flavoprotein complex; α-KG, alpha-ketoglutarate. (B, C) Heart mitochondria were mock-treated or treated with filter-sterilized Spn culture supernatants (∼50 µM H_2_O_2_, collected after 4 h growth). (B) Complex I-driven respiration (CIDR) and Complex II-driven respiration (CIIDR) were measured using Oroboros O2K FluoRespirometry with substrates: (CIDR) 10 mM pyruvate, 5 mM malate, 2 mM ADP, or 20 mM glutamate, 5 mM malate, 2 mM ADP; (CIIDR) 20 mM succinate, 2 mM ADP. (C) CIDR was measured with fatty acid substrates oleate and octanoate. (D–F) Enzymatic activities of aconitase, glutamate dehydrogenase (GDH), and α- ketoglutarate dehydrogenase were measured in heart mitochondria mock-treated or treated with Spn supernatants (∼50 µM H_2_O_2_), as described in Materials and Methods. (G) GDH activity was assessed in Calu-3 lung epithelial cells infected with Spn strains. (H, I) RNA-seq analysis of human bronchial Calu-3 cells mock-infected or infected with Spn strains. Heatmaps depict expression of (H) mitochondria-related or (I) oxidative stress-related genes; NS, not significant. (J) Diagram of cysteine oxidation experimental design. (K) Chemical reaction illustrating H_2_O_2_-mediated cysteine oxidation. (L) Cytosolic fraction appearance post-DCP-Rho1 incubation. (M, N) Western blot analysis of (M) cytosolic and (N) mitochondrial fractions using anti-TRITC antibodies (anti-RhoI), with β-actin and cytochrome c oxidase subunit II as respective loading controls. (B–G) Data represent mean ± SEM; one-way ANOVA with Dunnett’s post hoc test, ns, not significant; *p < 0.05; **p < 0.005; ***p < 0.0005; ****p < 0.0001.

β-oxidation of medium-chain and long-chain fatty acids in the mitochondrial matrix produces acetyl-CoA, via electron transfer flavoprotein (ETF, Fig. 4B), which reacts with oxaloacetate to form citrate in the TCA cycle. We assessed whether supplementing oleate (long-chain) or octanoate (medium-chain) to isolated mitochondria incubated with Spn cultures inhibits CIDR. Compared to the mock control, cultures of both strains significantly inhibited CIDR (Fig. 4C), further indicating that the inhibition induced by Spn cultures occurs at or downstream of CS.

In the mitochondria, aconitase (ACO2) is located downstream of CS and catalyzes the reversible isomerization of citrate to isocitrate. The resulting isocitrate is then decarboxylated to α-ketoglutarate by isocitrate dehydrogenase 3 (IDH3A) (Fig. 4A). Glutamate in the mitochondria is converted to α-ketoglutarate by glutamate dehydrogenase (GDH), which is then decarboxylated to succinyl-CoA by α-ketoglutarate dehydrogenase (OGDHC) (Fig. 4A). We assessed the activity of ACO2, GDH, and OGDHC in isolated mitochondria exposed to cultures of Spn strains producing H_2_O_2_, comparing them to mock-infected conditions and to cultures of Δ*spxB*Δ*lctO* mutants. Both TIGR4 and EF3030, caused a significant inhibition of the enzymatic activity of ACO2 (Fig. 4D), GDH (Fig. 4E), and OGDHC (Fig. 4F), compared to mock-infected cultures or cultures of isogenic Δ*spxB*Δ*lctO* mutants.

To elucidate the mechanism of inhibition, sulfenic acid (RSOH) formation, a marker of cysteine oxidation, was assessed in human bronchial epithelial Calu-3 cells using DCP-Rho1 (26, 50) (Fig. 4J), a dimedone-based fluorogenic probe specific for RSOH (50) (Fig. 4K). Cells infected with wild-type EF3030 exhibited significantly elevated RSOH levels in cytosolic (Fig. 4L, 4M), and mitochondrial (Fig. 4N) fractions compared to mock-infected cells or those infected with the H_2_O_2_-deficient Δ*spxB*Δ*lctO* mutant. These results indicate that Spn-H_2_O_2_ induces oxidative modification of cysteine residues in TCA cycle enzymes (ACO2, GDH, OGDHC), contributing to their inhibition. To further validate these findings in the same host cell model, we measured GDH activity in Calu-3 cells, either mock-infected or infected with Spn. Consistent with the *ex vivo* data, GDH activity in Spn-infected Calu-3 cells was significantly reduced compared to mock-infected controls or cells infected with Δ*spxB*Δ*lctO* (Fig. 4G), supporting the notion that Spn-H_2_O_2_ directly inhibits GDH, disrupting the TCA cycle and driving metabolic reprogramming in lung epithelial cells.

### Spn-mediated transcriptional regulation of mitochondrial genes in lung cells

RNA sequencing (RNAseq) analysis of bronchial Calu-3 cells, a differentiated lung epithelial model derived from adenocarcinoma, revealed extensive transcriptional reprogramming following infection with TIGR4. Compared to mock-infected controls, TIGR4 infection dysregulated 2,684 genes (FDR <0.05, log2 fold change >0.5), a response largely driven by SpxB, as infection with the isogenic TIGR4Δ*spxB* mutant dysregulated only 163 genes (Fig. S3A and S3B). Gene ontology (GO) analysis indicated that TIGR4 infection induces a broad stress response, with significant enrichment of pathways related to DNA damage response, cell cycle regulation, and mitochondrial function (Fig. S3C and S3D). In contrast, TIGR4Δ*spxB* infection elicited a narrower response, primarily involving NF-κB signaling and apoptosis pathways. Notably, TIGR4 infection dysregulated 237 mitochondrial-related genes (GO:0005739), compared to a few in TIGR4Δ*spxB*-infected cells, highlighting SpxB’s critical role in mitochondrial gene regulation (Table S1). Calu-3 cells, exhibiting a metabolic profile less dependent on Warburg metabolism than more aggressive cancer cell lines, provide a physiologically relevant model for studying epithelial responses to bacterial infection (51).

Among the dysregulated genes, TIGR4 infection significantly upregulated IDH3A (isocitrate dehydrogenase 3) and PDK1 (pyruvate dehydrogenase kinase 1) RNA expression in Calu-3 cells (Fig. 4H), while downregulating PC (pyruvate carboxylase), PDK2 (pyruvate dehydrogenase kinase 2), and PDPR (pyruvate dehydrogenase phosphatase regulatory subunit). The proteins encoded by these genes regulate the pyruvate dehydrogenase complex (PDC), which controls pyruvate entry into the mitochondrial TCA cycle. Upregulation of PDK1 likely inhibits PDC activity, while concurrent downregulation of PDK2 and PDPR further suppresses PDC function, potentially limiting mitochondrial oxidative phosphorylation and mitigating oxidative stress. This pattern was absent in TIGR4Δ*spxB*-infected cells, which lack pyruvate oxidase and H_2_O_2_ production. However, a similar dysregulation was observed with the H_2_O_2_-producing TIGR4Δ*lctO* mutant (Fig. 4H), emphasizing SpxB-mediated oxidative stress in driving these changes.

Supporting SpxB-driven transcriptional reprogramming, genes encoding oxidative stress response enzymes, including SOD2 (superoxide dismutase 2), TXNRD1 (thioredoxin reductase 1), HMOX1 (heme oxygenase 1), and NFKB1 (nuclear factor kappa B subunit 1), were upregulated in Calu-3 cells infected with TIGR4 and TIGR4Δ*lctO* (Fig. 4I). Expression remained unchanged in TIGR4Δ*spxB*-infected cells, highlighting SpxB’s role in eliciting a robust antioxidant defense. Similar dysregulation of PDK1 and SOD genes was observed in non-cancerous bronchial BEAS-2B cells infected with Spn strain D39 (52) (explained later, Fig. 7E).

### Spn-H_2_O_2_-driven metabolic reprogramming in lung epithelial cells

Inhibition of key TCA cycle enzymes, including ACO2, and OGDHC, by Spn may lead to citrate accumulation in host cells. To investigate this, intracellular and extracellular citrate concentrations were measured in mock-infected Calu-3 bronchial epithelial cells and compared to cells infected with Spn strains TIGR4 or TIGR4Δ*spxB*Δ*lctO* for 8 h. Extracellular citrate levels, quantified by high- performance liquid chromatography (HPLC), were significantly elevated in TIGR4- and TIGR4Δ*spxB*Δ*lctO*-infected cells (median 227 µM and 441 µM, respectively) compared to mock-infected controls (4.04 µM) (Fig. 5A). Citrate levels in Spn culture supernatants (without host cells) were comparable between TIGR4 and TIGR4Δ*spxB*Δ*lctO*, suggesting bacterial contribution to extracellular citrate (Fig. 5A). However, intracellular citrate concentrations were significantly higher in TIGR4-infected cells compared to mock-infected or TIGR4Δ*spxB*Δ*lctO*- infected cells (Fig. 5B), indicating that H_2_O_2_-producing Spn exacerbates mitochondrial citrate accumulation, likely due to TCA cycle inhibition. This citrate accumulation in TIGR4-infected Calu-3 cells aligns with findings by Surabhi et al. (2021)(49), confirming that Spn-H_2_O_2_- mediated TCA cycle inhibition is a key driver of metabolic reprogramming in infected host cells. Metabolite analysis revealed that Spn infection enhances glucose consumption.

**Figure 5.**
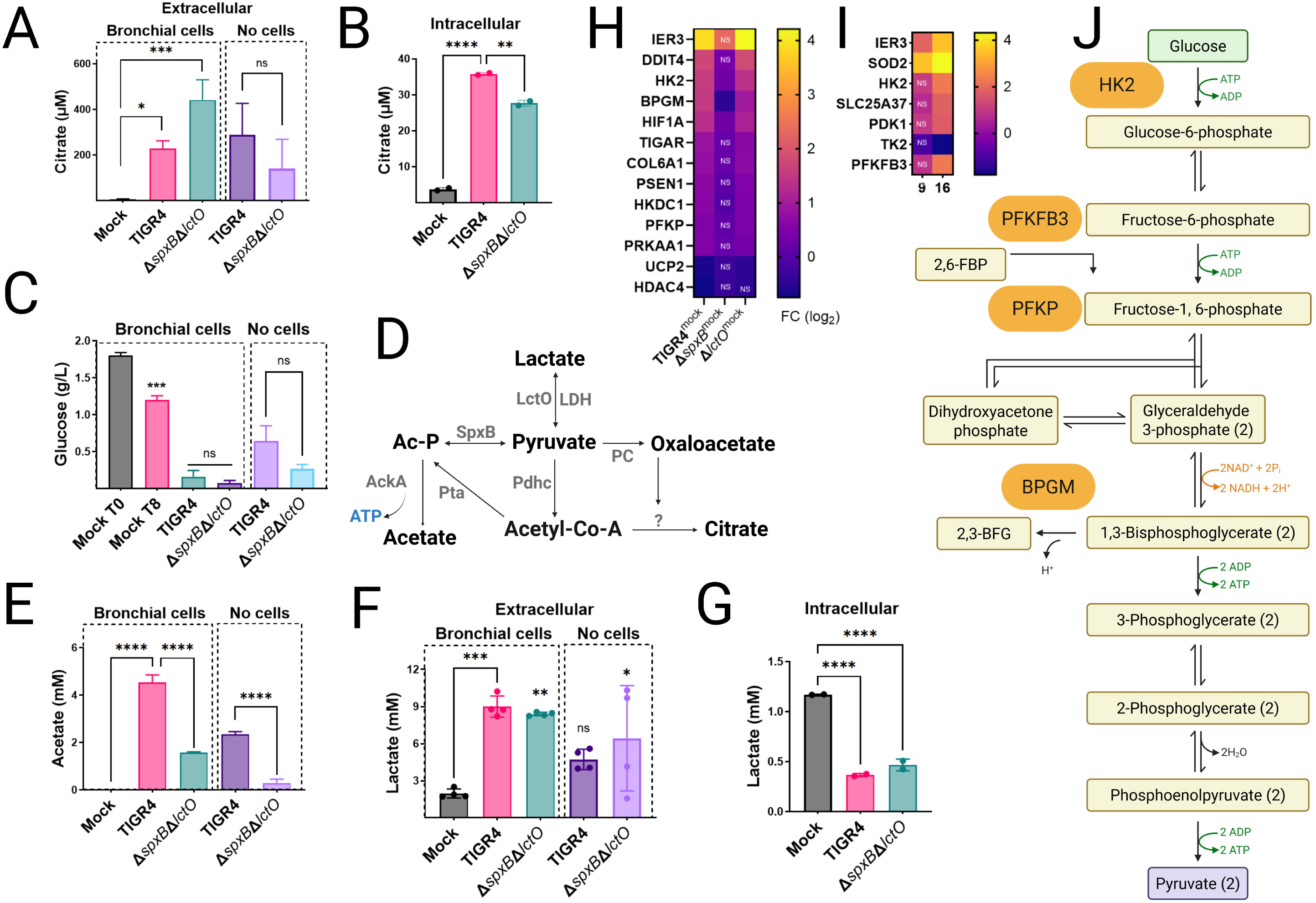
Spn-H_2_O_2_ drives Warburg-like metabolic reprogramming in lung epithelial cells. (A–C, E–G) Human bronchial Calu-3 cells were mock-infected, infected with Spn strains, or incubated with Spn cultured in cell culture medium for 8 h. Culture supernatants (extracellular) or cell pellets (intracellular) were harvested to quantify levels of (A, B) citrate, (C) glucose, (E) acetate, and (F, G) lactate, as described in Materials and Methods. Data represent mean ± SEM; one-way ANOVA with Dunnett’s post hoc test (versus mock-infected cells) or unpaired Student’s t-test (versus TIGR4 strain); ns, not significant; *p < 0.05; **p < 0.005; ***p < 0.0005; ****p < 0.0001. (D) Diagram of pyruvate node reactions under aerobic conditions, incorporating oxaloacetate and citrate. Enzymes are highlighted in gray. (H) RNA-seq analysis of Calu-3 cells or (I) BEAS 2B mock-infected or infected with Spn strains. Heatmap depicts expression of glycolysis-related genes; NS, not significant. (K) Schematic of the canonical glycolysis pathway, highlighting genes encoding enzymes upregulated in RNA-seq analysis.

Baseline glucose in Calu-3 cells was 1.8 g/L, decreasing to 1.2 g/L in mock-infected cells after 8 h. In TIGR4- and TIGR4Δ*spxB*Δ*lctO*-infected cells, glucose levels dropped below 0.16 g/L (Fig. 5C), while Spn cultures alone consumed less than 0.65 g/L, indicating a host-specific response likely driven by SpxB-mediated oxidative stress and TCA cycle suppression. Acetate production, absent in mock-infected cells, increased to 4.54 mM in TIGR4-infected cells but was lower in TIGR4Δ*spxB*Δ*lctO*-infected cells (1.58 mM) (Fig. 5E). Spn cultured alone produced 2.35 mM (TIGR4) and 0.29 mM (TIGR4Δ*spxB*Δ*lctO*) acetate, reflecting SpxB’s role in acetate synthesis via acetyl phosphate under aerobic conditions (Fig. 5D). Lactate levels further highlighted metabolic shifts. Extracellular lactate in mock-infected cells was 1.96 mM, increasing to 8.9 mM (TIGR4) and 8.4 mM (TIGR4Δ*spxB*Δ*lctO*) in infected cells, with Spn alone producing 4.7 mM (TIGR4) and 6.3 mM (TIGR4Δ*spxB*Δ*lctO*) (Fig. 5F). Intracellular lactate, 1.17 mM in mock-infected cells, decreased significantly in infected cells, likely due to Spn-induced cytotoxicity (Fig. 5G). RNA-seq analysis revealed significant upregulation of glycolysis-related genes in TIGR4-infected cells, including IER3, DDIT4, HK2, HIF1A, and PFKP (log_2_ fold change 1–4, FDR <0.05), with IER3, HK2 and PFKP also upregulated in TIGR4Δ*lctO*-infected cells (Fig. 5H, 5J). IER3 (immediate early response 3) and HIF1A (hypoxia-inducible factor 1-alpha) transcriptionally regulates glycolytic genes glycolytic genes promoting a Warburg-like glycolytic shift in cancer cells (53, 54). Similarly, RNA-seq analysis of BEAS-2B bronchial epithelial cells infected with Spn for 9 or 16 h revealed upregulated expression of glycolysis-related genes HK2, PDK1, TK2, PFKFB3, and IER3 (Fig. 5I). These findings in a non-cancerous cell line corroborate the Warburg-like glycolytic shift observed in Calu-3 cells. In TIGR4Δ*spxB*-infected cells, most genes were unchanged except for HK2, BPGM, and HIF1A, underscoring SpxB-H_2_O_2_’s role in glycolytic reprogramming.

Collectively, Spn-H_2_O_2_ induces profound metabolic reprogramming in Calu-3 cells, marked by citrate accumulation, enhanced glucose consumption, increased extracellular lactate and acetate, and selective upregulation of glycolytic genes, reflecting a host response to SpxB-mediated mitochondrial impairment.

### Infection with *S. pneumoniae* does not cause loss of alveolar mitochondrial membrane potential

Increased permeability of the mitochondrial inner membrane (IMM) can dissipate the proton electrochemical gradient and destabilize mitochondrial membrane potential (ΔΨm), both of which are essential for ATP synthesis via ATP synthase. Given that Spn-H_2_O_2_ inhibits the TCA cycle, we hypothesized that enhanced mitochondrial permeability contributes to this effect. To test this, ΔΨm—established primarily by proton translocation through complexes I, III, and IV of the electron transport chain (ETC)—was assessed, as it, together with the proton gradient, constitutes the proton-motive force driving ATP production (55). In healthy mitochondria, ΔΨm is tightly regulated; its dissipation or reduction signals impaired ATP synthesis, mitochondrial dysfunction, and increased reactive oxygen species (ROS) production. Here, we evaluated the impact of Spn-derived H_2_O_2_ on ΔΨm in A549 alveolar epithelial cells using flow cytometry with the potentiometric dye JC-1.

In cells with intact mitochondrial function, the cell-permeant dye JC-1 accumulates in polarized mitochondria, forming aggregates that exhibit a balanced fluorescence emission profile, with an approximately equal ratio of Texas Red (∼590 nm) to FITC (∼530 nm). This dual-emission state is detected as a +Texas Red/+FITC double-positive population in flow cytometry, indicative of robust ΔΨm. Consistent with this, uninfected A549 cells stained with JC-1 displayed this characteristic pattern, reflecting maintained ΔΨm (Fig. 6A). In contrast, treatment with carbonyl cyanide m-chlorophenyl hydrazone (CCCP), a known ΔΨm uncoupler, induced complete depolarization, shifting the population to -Texas Red/+FITC single-positive cells (Fig. 6B).

**Figure 6.**
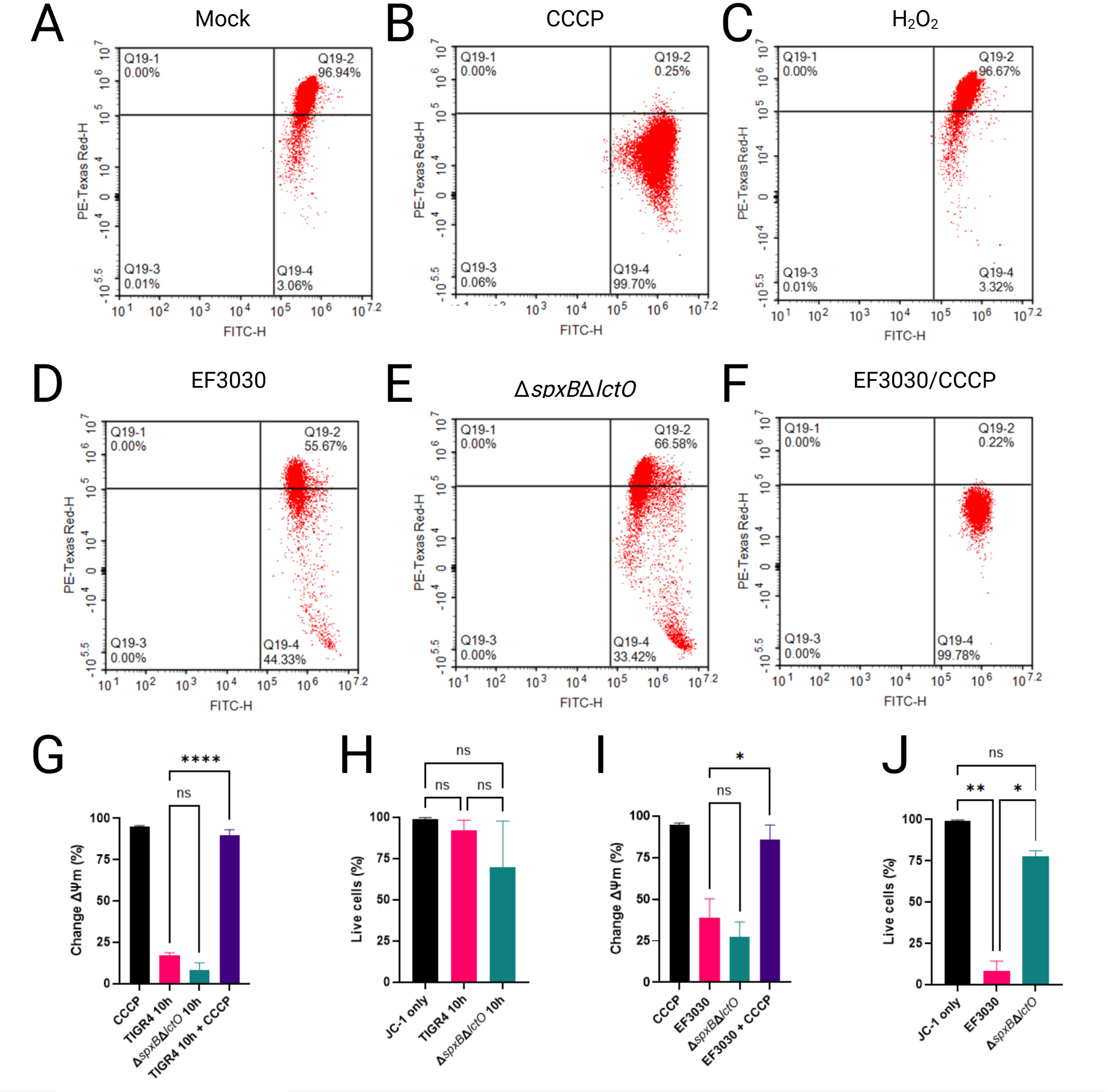
*Streptococcus pneumoniae*-derived. **H**_2_**O**_2_ **does not significantly alter mitochondrial membrane potential in lung epithelial cells.** (A–H) Human alveolar A549 cells were mock-treated, treated with CCCP, H_2_O_2_, EF3030, EF3030Δ*spxB*Δ*lctO* or EF3030 and CCCP for 10 h. Cells were stained with JC-1 dye for flow cytometric analysis and counterstained with SYTOX Blue to exclude dead cells. (A–F) Scatter plots showing JC-1 fluorescence (PE-TexasRed/FITC) for each condition, with gating to identify live cells. (G and I) Quantification of change in mitochondrial membrane potential (Δψm) or (H and J) percent of live cells expressed as mean ± SEM from three independent experiment; one-way ANOVA with Dunnett’s post hoc test; ns, not significant; *p < 0.05; **p < 0.01; ****p < 0.0001.

To assess the effects of Spn-derived H_2_O_2_ on mitochondrial function, A549 cells were infected with wild-type strains TIGR4 and EF3030, or their respective H_2_O_2_-deficient mutants TIGR4Δ*spxB*Δ*lctO* and EF3030Δ*spxB*Δ*lctO*. ΔΨm was then quantified using JC-1 staining and flow cytometry. Relative to the CCCP-treated positive control, which induces complete ΔΨm dissipation by enhancing proton permeability across the IMM, infections with TIGR4 and EF3030 resulted in significantly less mitochondrial depolarization (Figs. 6D, 6G, 6I). Similarly, treatment of A549 cells with 2 mM H_2_O_2_ alone did not alter ΔΨm (Fig. 6C), suggesting that H_2_O_2_ at this concentration does not directly impair mitochondrial membrane integrity. Furthermore, A549 cells infected with TIGR4 or EF3030 and subsequently treated with CCCP exhibited complete ΔΨm dissipation (Figs. 6F, 6G, 6I), confirming that lung epithelial cells maintain an intact ΔΨm despite Spn infection over this timeframe. No significant changes in ΔΨm were observed in cells infected with TIGR4Δ*spxB*Δ*lctO* or EF3030Δ*spxB*Δ*lctO* compared to their wild-type counterparts, indicating that Spn-derived H_2_O_2_ does not substantially affect ΔΨm under these conditions.

To evaluate cytotoxicity, the same population of infected A549 cells was analyzed using SYTOX Blue staining and flow cytometry. Compared to mock-infected controls, no significant reduction in cell viability was observed in A549 cells infected with TIGR4 or its H_2_O_2_-deficient mutant TIGR4Δ*spxB*ΔlctO (Fig. 6H). In contrast, infection with EF3030 resulted in a significant decrease in cell viability relative to mock-infected controls and the corresponding H_2_O_2_- deficient mutant EF3030Δ*spxB*Δ*lctO* (Fig. 6J), likely due to elevated H_2_O_2_ production by EF3030 relative to TIGR4 (Fig. 3A). Collectively, these results suggest that Spn infection does not substantially impair ΔΨm, consistent with H_2_O_2_ targeting TCA cycle enzymes (e.g., ACO2, OGDHC) rather than ETC complexes, thereby preserving mitochondrial membrane integrity over the experimental timeframe.

### Spn-Derived H_2_O_2_ modulates limited alveolar apoptosis via TCA cycle Inhibition

Having demonstrated that Spn-derived H_2_O_2_ inhibits the tricarboxylic acid (TCA) cycle without disrupting mitochondrial membrane potential (ΔΨm) in lung epithelial cells, we explored whether this metabolic perturbation triggers cell death by impairing oxidative phosphorylation (OXPHOS). Apoptosis was evaluated using the terminal deoxynucleotidyl transferase biotin- dUTP nick end labeling (TUNEL) assay, which detects DNA fragmentation—a characteristic of late-stage apoptosis—by fluorescently labeling 3’-OH termini in fragmented DNA, facilitating quantitative analysis of apoptotic cell populations via confocal microscopy. Additionally, activation of caspase-3 and caspase-7 was assessed using flow cytometry and confocal microscopy. Mock-infected A549 alveolar epithelial cells displayed minimal DNA fragmentation, whereas DNase I-treated cells, serving as a positive control, exhibited a marked increase in fluorescence intensity, predominantly localized to nuclear regions (Fig. 7A, Fig. S4A), validating the assay. Cells treated with staurosporine, a known apoptosis inducer, showed reduced viability, with ∼60% of viable cells exhibiting activated caspase-3/7 (Fig. 7C). Three- dimensional reconstruction of z-stacks, including XY, XZ, and YZ optical sections (Fig. 7E, Fig. S5), along with time-lapse microscopy (Video S1), confirmed apoptosis induction in staurosporine-treated cells.

**Figure 7.**
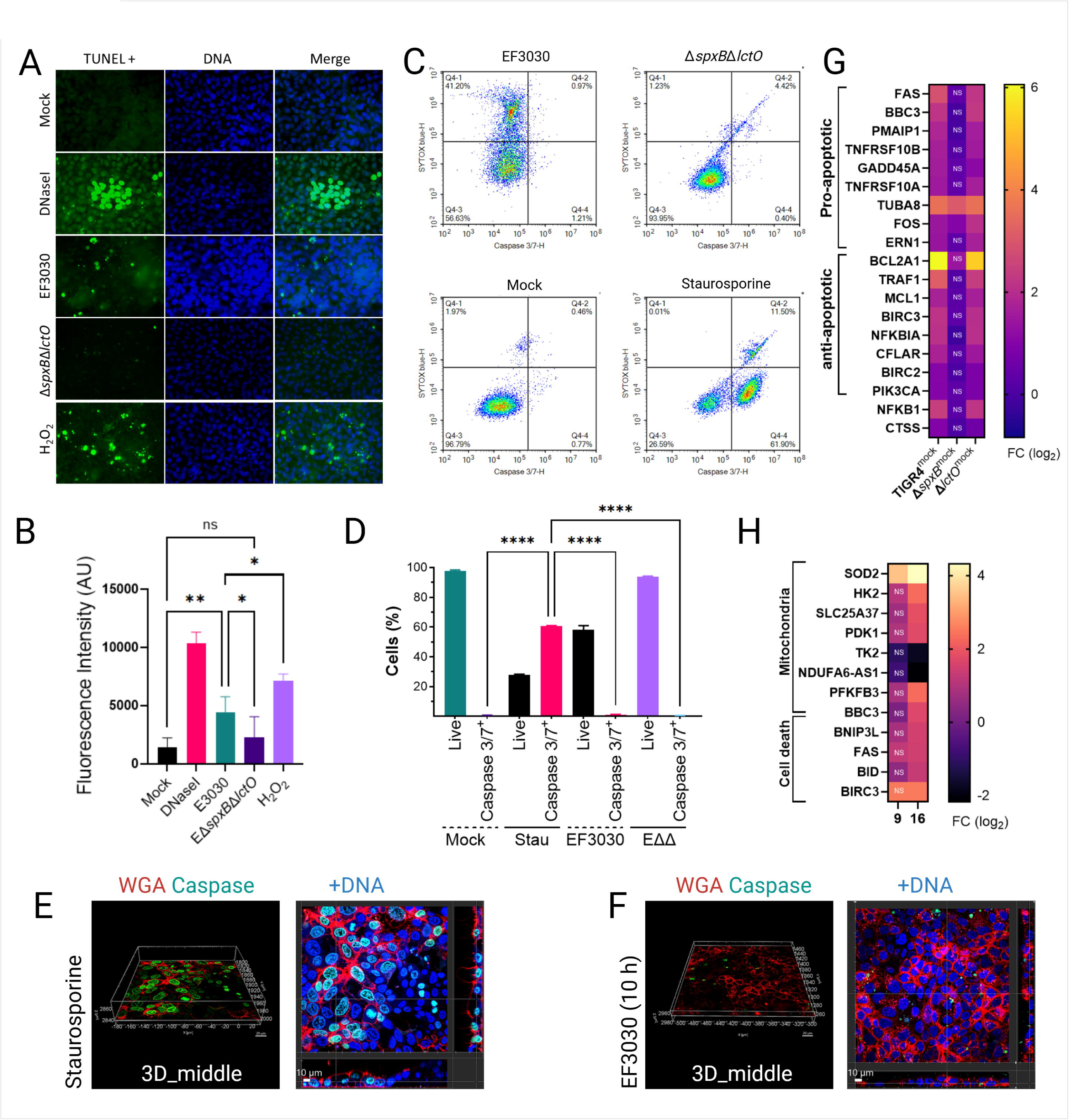
Limited contribution of Spn-H_2_O_2_ to apoptosis in lung epithelial cells. (A) Human alveolar A549 cells were mock-infected, treated with DNaseI (1U), treated with H_2_O_2_ (2 mM) added every 2 h, or infected with TIGR4, TIGR4Δ*spxB*Δ*lctO*, EF3030, or EF3030Δ*spxB*Δ*lctO* for 10 h. Cells were stained with the Click-iT Plus TUNEL assay kit, and mounted on slides with DAPI-containing resin and analyzed by confocal microscopy using the Imaris software. (B) Fluorescence intensity (FITC/GFP) from each condition, reported as arbitrary units (AU), quantifying TUNEL staining. (C) A549 cells were mock-infected or infected with EF3030, EF3030Δ*spxB*Δ*lctO* for 10 h or treated with staurosporine (10 µM, 4 h), then stained with Cell Event Caspase-3/7 Green Detection Reagent (2 µM) and SYTOX Blue Dead Cell Stain (1 µM) prior to flow cytometric analysis. Representative bivariate density plots show the percentages of single A549 cells within the P3 gate: non-apoptotic SYTOX blue–positive cells (Q4-1), SYTOX blue–negative non-apoptotic cells (Q4-3), and caspase-3/7–positive, SYTOX blue–negative cells (Q4-4) for each condition. (D) Percentage of SYTOX Blue-negative live cells, and among those, the proportion of caspase-3/7 positive cells. (B, D) Data represent mean ± SEM from two independent experiments with four (B) or two (D) internal replicates each; one-way ANOVA with Dunnett’s post hoc test; ns, not significant; *p < 0.05; **p < 0.01; ****p < 0.0001. (E-F) Cells treated with staurosporine (10 µM) for 7 hours or infected with EF3030 for 10 hours were stained with WGA, caspase-3/7 reagent, and DAPI as indicated. Z- stacks were acquired in real-time using confocal microscopy at the specified time points. Displayed are the middle sections of 3D images from z-stacks (left panels) and the corresponding XY, XZ, and YZ optical sections (right panels). (D, E) RNA-seq analysis of Calu- 3 (D) or BEAS-2B (E) cells mock-infected or infected for 9 or 16 h with Spn strains. Heatmaps depict expression of mitochondria-related and apoptosis-related genes

A549 cells were infected for 10 h with wild-type Spn strains TIGR4 and EF3030, or their H_2_O_2_-deficient mutants TIGR4Δ*spxB*Δ*lctO* and EF3030Δ*spxB*Δ*lctO*. TUNEL-positive cells were observed infrequently across all infected conditions (Fig. 7A, Fig. S4), though the signal in infected cells was significantly higher than in mock-infected controls (Fig. 7B, Fig. S4B), yet markedly lower than in DNase I-treated cells. Notably, cells infected with H_2_O_2_-producing strains TIGR4 and EF3030 exhibited a significantly greater TUNEL signal compared to those infected with H_2_O_2_-deficient mutants (Fig. 7B, Fig. S4B), indicating that Spn-derived H_2_O_2_ promotes cell death. Consistent with this, flow cytometry analyses revealed a significantly greater proportion of viable cells in infections with H_2_O_2_-deficient mutants compared to their wild-type counterparts (Fig. 7D). However, activation of caspase-3/7 in Spn-infected cells was not detected at 10 h post-infection, as confirmed by flow cytometry, confocal microscopy and time-lapse studies (Fig. 7C, 7F, Fig. S4C, Fig. S5 and Video S1). These findings suggest that H_2_O_2_-mediated inhibition of TCA cycle enzymes (e.g., ACO2, OGDHC), which supply electron donors for OXPHOS, contributes to a limited apoptotic response, consistent with the minimal ΔΨm disruption observed previously.

To further explore the role of H_2_O_2_ in apoptosis, A549 cells were treated with 2 mM H_2_O_2_ every 2 h for 8 h, simulating the delayed accumulation of Spn-derived H_2_O_2_ during infection, which becomes detectable after 4 hours as cellular substrates are depleted (46). TUNEL analysis revealed that the apoptotic signal in H_2_O_2_-treated cells was comparable to that in wild-type Spn-infected cells but significantly lower than in DNase I-treated controls (Fig. 7A, 7B). These results confirm that Spn-derived H_2_O_2_ contributes to limited alveolar cell apoptosis, likely via metabolic stress from TCA cycle inhibition, though its effect is less pronounced than that of canonical apoptosis inducers.

Gene expression analysis revealed a complex interplay of apoptosis regulation in human bronchial cells infected with H_2_O_2_-producing strains TIGR4 and TIGR4Δ*lctO* compared to mock-infected controls and the H_2_O_2_-deficient mutant TIGR4Δ*spxB* (Fig. 7G). Pro-apoptotic genes, including FAS and BBC3, were significantly upregulated in TIGR4-infected cells (log_2_ fold-change [FC] 1–6; FDR < 0.05), promoting apoptosis via death receptor signaling and stress response pathways. Concurrently, anti-apoptotic genes such as BCL2A1 (BCL2-family member, prevent Cyt*c* release) and TRAF1 were also upregulated, likely counteracting cell death by inhibiting caspase activation and enhancing survival through NF-kappaB signaling (NFKB1). This dual upregulation of pro- and anti-apoptotic genes, absent in TIGR4Δ*spxB*- infected cells, likely underlies the limited apoptosis observed in the TUNEL assay, as the balance between these opposing pathways mitigates extensive cell death despite *spxB*- mediated H_2_O_2_ stress. Similar dual regulation of pro- and anti-apoptotic genes was observed in BEAS-2B cells infected with Spn reference strain D39 for 16 hours (Fig. 7H), mirroring patterns reported in MYC-driven cancer biology (56). These findings highlight a novel mechanism by which Spn modulates host cell survival to sustain infection.

## Discussion

This study unveils a novel and unique mechanism by which Spn disrupts host cell metabolism through H_2_O_2_-mediated inhibition of the TCA cycle, coupled with extensive transcriptional and metabolic reprogramming in lung epithelial cells. To our knowledge, direct inhibition of the TCA cycle enzymes by a bacterial pathogen has not been described prior to this work, positioning Spn’s H_2_O_2_-driven metabolic disruption as a groundbreaking insight into pneumococcal pathogenesis. Spn-derived H_2_O_2_, primarily produced by pyruvate oxidase (SpxB), targets key mitochondrial enzymes—aconitase (ACO2), glutamate dehydrogenase (GDH), and α-ketoglutarate dehydrogenase (OGDHC)—significantly inhibiting their activities. H_2_O_2_-induced cysteine oxidation, marked by elevated sulfenic acid levels in EF3030-infected Calu-3 cells, drives this inhibition. This leads to citrate accumulation, as evidenced by elevated intracellular and extracellular citrate levels in TIGR4-infected Calu-3 cells compared to mock-infected controls or cells infected with the H_2_O_2_-deficient mutant TIGR4Δ*spxB*Δ*lctO*. This H_2_O_2_- driven citrate accumulation, also observed by Surabhi et al. (2021)(49), highlights Spn’s unique strategy to disrupt host metabolism via TCA cycle inhibition

This metabolic bottleneck downstream of citrate synthase impairs NADH production and succinate reduction, critical electron donors for OXPHOS, aligning with the observed inhibition of CIDR at H_2_O_2_-concentrations as low as 5 µM. In contrast, CIIDR, which utilizes substrates acting downstream of the enzymes inhibited by Spn-H_2_O_2_, required higher concentrations (25 µM) for inhibition. Catalase rescue and exogenous H_2_O_2_ treatment assays (Fig. 3G–3J) revealed that Spn-H_2_O_2_-mediated inhibition of CIDR is reversible and redox-sensitive, suggesting dynamic modulation of mitochondrial function. This reversibility highlights potential therapeutic strategies to mitigate Spn-induced metabolic stress by targeting redox pathways. Notably, the absence of significant changes in ETC complex activities (Fig. 3K, Fig. S2) underscores that Spn-H_2_O_2_ targets upstream TCA cycle enzymes rather than the ETC itself, a mechanism distinct from the ETC decline associated with human diseases like neurodegeneration, cancer, and diabetes (57).

The hallmark of pneumococcal disease is Spn’s ability to invade host cells and manipulate their metabolism to establish infection (58). Our findings demonstrate that H_2_O_2_- mediated inhibition of the TCA cycle disrupts host energy metabolism and induces transcriptional reprogramming, shifting lung epithelial cells toward a Warburg-like phenotype(59) that fosters bacterial proliferation. The NAD^+^ salvage pathway, which supplies electron donors for host antibacterial responses against Spn (52), is likely impaired by reduced NADH production resulting from TCA cycle suppression. This metabolic disruption compromises host defenses, enabling Spn to exploit host resources, as evidenced by significant glucose depletion in TIGR4-infected Calu-3 cells. Furthermore, transcriptional reprogramming in TIGR4-infected cells amplifies this metabolic shift. Upregulation of glycolysis-associated genes (HK2, PFKP, HIF1A) and downregulation of pyruvate dehydrogenase complex regulators (PC, PDK2, PDPR) indicate the Warburg-like metabolic switch toward glycolysis (59), likely a compensatory response to H_2_O_2_-induced mitochondrial dysfunction. To further validate the Warburg-like metabolic shift in a non-cancerous context, RNA-seq analysis of BEAS-2B bronchial epithelial cells infected with Spn strain D39 (Fig. 5I) confirmed upregulated expression of glycolysis-related genes (HK2, PDK1, TK2, PFKFB3) and IER3, reinforcing the physiological relevance of these transcriptional changes across cell types (52). This reprogramming, absent in TIGR4Δ*spxB*Δ*lctO*-infected cells, underscores the critical role of SpxB in driving host metabolic stress.

Spn- H_2_O_2_ induces mitochondrial aggregation and apical migration in A549 cells (Fig. 2B–C) and Calu-3 cells (Fig. S1), indicative of disrupted mitochondrial dynamics that amplify oxidative stress. The interaction between the cytoskeleton and the mitochondrial outer membrane is critical for regulating mitochondrial respiration (60). Mutations in microtubule-associated proteins, such as plectin or desmin, impair mitochondrial network integrity, and similarly, Spn-H_2_O_2_-mediated microtubule disruption, absent in H_2_O_2_-deficient mutants (17, 23), likely drives mitochondrial aggregation and mislocalization in lung epithelial cells, potentially contributing to metabolic dysfunction. Supporting this, the absence of βII-tubulin in cardiomyocytes correlates with a shift to glycolysis-dependent metabolism (Warbug effect) (61). The increased extracellular lactate and acetate in TIGR4-infected cells (Fig. 5E–F) reflect Spn’s manipulation of host glycolytic flux, potentially fueling bacterial metabolism while stressing host cells. Notably, mitochondrial membrane potential (ΔΨm) remains intact in TIGR4- and EF3030-infected cells, indicating that H_2_O_2_ targets TCA cycle enzymes rather than ETC complexes or membrane integrity. The preservation of mitochondrial membrane potential (ΔΨm) in TIGR4- and EF3030-infected cells (Fig. 6) reflects the selective inhibition of TCA cycle enzymes by Spn-H_2_O_2_, rather than ETC damage, as exogenous H_2_O_2_ might cause (62). This distinction underscores the physiological relevance of our infection model, where H_2_O_2_ levels (499.5–1,119 µM) target upstream metabolic pathways. This preservation of ΔΨm aligns with the limited apoptosis observed in TUNEL assays, where TIGR4- and EF3030-infected cells showed significantly higher apoptotic signals than H_2_O_2_-deficient mutants but far less than DNase I-treated controls. The balanced upregulation of pro-apoptotic (FAS, BBC3) and anti- apoptotic (BCL2A1, TRAF1) genes in TIGR4-infected cells likely mitigates extensive cell death, enabling Spn to maintain host cell viability for sustained infection.

The decline in ETC activity is a hallmark of many human diseases, yet our study uniquely demonstrates that Spn directly targets the TCA cycle, upstream of the ETC, to disrupt host metabolism. The oxidation of soluble Cyt*c* by Spn-H_2_O_2_, forming tyrosyl radicals, further impairs mitochondrial function by disrupting electron transfer, amplifying ROS production, and exacerbating metabolic stress. This mechanism not only enhances Spn’s virulence but also parallels metabolic disruptions, suggesting broader implications for understanding host- pathogen interactions. Elevated intracellular ROS levels in lung epithelial cells infected with wild-type Spn strains TIGR4 and EF3030, compared to Δ*spxB*Δ*lctO* mutants or mock-infected controls, support a mechanistic link between extracellular tyrosyl radical formation and intracellular oxidative stress, amplifying mitochondrial dysfunction (45, 46).

While heart mitochondria provided a robust model for studying Spn-induced mitochondrial dysfunction in this study, their physiological differences from lung epithelial mitochondria, including substrate preference (fatty acids vs. glucose/glutamine) and higher respiratory capacity, may limit direct extrapolation to the lung context. These differences, coupled with distinct ROS handling capacities in lung epithelial mitochondria, suggest that tissue-specific responses to Spn-H_2_O_2_ may vary. Future studies using mitochondria isolated from lung epithelial cells could validate our findings and elucidate tissue-specific metabolic responses to Spn infection, enhancing the translational relevance of our results.

In conclusion, Spn employs H_2_O_2_-mediated inhibition of the TCA cycle as a novel strategy to reprogram host cell metabolism and the transcriptional profiles, thereby facilitating bacterial invasion and persistence. By disrupting NADH production and inducing glycolytic reprogramming, Spn impairs host antibacterial responses, including those dependent on the NAD^+^ salvage pathway(52), while enhancing its own survival. These findings underscore the therapeutic potential of targeting SpxB activity, such as through SpxB inhibitors, or restoring TCA cycle function to attenuate pneumococcal disease severity. Recent evidence suggests that enhancing host cell NAD^+^ metabolism can suppress Spn replication (52). Future studies should delineate the molecular interactions between H_2_O_2_ and TCA cycle enzymes, investigate the role of host antioxidant defenses in mitigating Spn-induced metabolic stress, and employ *in vivo* models to elucidate how Spn’s metabolic manipulation drives disease progression. Such investigations may inform the development of innovative therapeutic strategies for pneumococcal infections.

## Material and Methods

### Bacterial strains, growth conditions, and preparation of inoculum

Spn strains utilized in this study are listed in Table 1. Strains were cultured on blood agar plates (BAP) containing 5% sheep’s blood from frozen bacterial stocks stored in skim milk, tryptone, glucose, and glycerin (STGG)(63) or in Todd-Hewitt broth supplemented with 0.5% (w/v) yeast extract (THY) and incubated at 37°C with 5% CO_2_. Inoculum for experiments was prepared as previously described (9). Overnight cultures of Spn on BAP were harvested in phosphate buffered saline (PBS) (pH 7.4), the OD_600_ of this suspension was obtained and adjusted to a final 0.1 density that contained ∼5.15 x 10^8^ cfu/ml.

**Table 1.** Strains utilized in this study.

| Strain | Description | Reference or Source |
| --- | --- | --- |
| TIGR4 | Invasive clinical isolate, capsular serotype 4, sensitive to antibiotics | (36) |
| TIGR4ΔspxB (SPJV29) | TIGR4 derivative with insertionally inactivated spxB::ermB, erythromycin resistance, hydrogen peroxide deficient strain | (37) |
| TIGR4ΔlctO (SPJV42) | TIGR4 derivative with insertionally inactivated lctO::aad9, spectinomycin resistance, hydrogen peroxide deficient strain | (9) |
| TIGR4ΔspxBΔlctO (SPJV41) | TIGR4 derivative with spxB and lctO gene deletion by insertion-deletion with erythromycin and spectinomycin cassette, respectively, hydrogen peroxide deficient strain | (9) |
| EF3030 | Invasive clinical isolate, capsular serotype 19F, sensitive to antibiotics | (69) |
| EF3030ΔspxBΔlctO (SPJV49) | EF3030 derivative with spxB and lctO gene deletion by insertion-deletion with erythromycin and spectinomycin cassette, respectively, hydrogen peroxide deficient strain | (37) |
| R6 | Laboratory unencapsulated isolate, derivative of D39, serotype 2 | (35) |

### Cell cultures

A549 human type II alveolar epithelial cells (ATCC, CCL-185) were cultured in Dulbecco’s Modified Eagle’s Medium (DMEM) (Gibco) supplemented with 10% fetal bovine serum (FBS), 2 mM L-glutamine (Gibco), and 100 U/mL of penicillin-streptomycin (Gibco) in 25 cm^2^ flasks (Fisher). Calu-3 cells (ATCC, HTB-55) were cultured in Eagle’s Minimum Essential Medium (EMEM, ATCC) supplemented with 10% FBS and 100 U/mL of penicillin-streptomycin (Gibco). Cell passages were performed by trypsinization with Trypsin-EDTA (0.25%), phenol red (Gibco). Cells were incubated at 37°C with 5% CO_2_ and were supplemented with fresh medium 2-3 times weekly and passaged to a new flask weekly or when cells reached ∼100% confluency.

### Infection of human bronchial and alveolar cells with *S. pneumoniae* strains

Cell infection experiments were performed using cells cultured for 8-10 days in the specified device. Cells were grown in 6-well plates with glass cover slips (Genesee, Fisher) or 8-well glass slides (Thermo Fisher Scientific, Nunc Lab-Tek II CC2) at 37°C with 5% CO_2_. Prior to infection, cells were washed three times with sterile PBS and then incubated in infection media consisting of DMEM or EMEM, supplemented with 5% FBS, 2 mM L-glutamine (only DMEM), and 1X HEPES (Gibco), referred as to infection medium. Cells were infected with Spn strains, mutant derivatives, or along with appropriate controls, and incubated at 37°C with 5% CO_2_ for various time points.

### Cytochrome c Oxidation Assay

To analyze the oxidation and degradation of heme in cytochrome c (Cytc) by Spn-H_2_O_2_, THY supplemented with 56 µM Cytc from horse heart (Sigma) was inoculated with Spn and incubated for 6 hours at 37°C with 5% CO2 in a 6-well plate. Supernatants were collected, centrifuged at 4°C for 5 minutes at 21,130×g, and transferred to a 96-well plate. The plate was then analyzed by spectroscopy, spanning wavelengths from 200 nm to 1000 nm, using an SYN H1 Hybrid Biotek reader (Agilent). As a control, 500 units (U) of catalase were added to a well containing THY-Cytc and Spn.

### Detection of radicals by spin-trapping

DEPMPO [5-(Diethoxyphosphoryl)-5-methyl-1- pyrroline-N-oxide, 98%, Cayman Chemical, P/N 10006435] was utilized to detect and identify the formation of radical species. Spn was cultured in THY and incubated for 4 h at 37°C. As a control, uninfected THY was incubated in the same condition. After incubation, the cell suspension was micro-centrifuged, and 32 μL of supernatant was used for each spin-trapping sample. The final reaction mixture contained, when added, 30 mM DEPMPO and 56 μM Cyt*c*, in a volume of 40 μL. Spn-H_2_O_2_ free sample was obtained by pretreating the bacterial supernatant liquid with 100 U catalase for 1 minute. Methods of radical detection are performed as described previously(5).

### Fluorescent staining of alveolar cells

A549 cells, or Calu-3 cells, seeded in a 6-well plate with glass coverslips were washed and added with infection media as mentioned above. Cells were then infected with Spn and incubated for 4, 6, or 8 h. After incubation with Spn, cells were fixed with 2% paraformaldehyde, permeabilized with 0.5% Triton X100, and blocked with 2% bovine serum albumin (BSA) following a previously described protocol(5). Post-blocking, cellular mitochondria were stained with an anti-mitochondria MTC02 monoclonal antibody (Invitrogen) at a concentration of 2 μg/mL in 2% BSA for 1 h. Cells were then washed with PBS and incubated for 1 h with a secondary goat anti-mouse FITC (SouthernBiotech) at a concentration of 20 μg/mL and phalloidin (7 units), or WGA-A555 at 1 μg/mL (ThermoFisher), in 2% BSA for 1 h. Cells were then washed, the coverslips were removed from the wells, allowed to dry, mounted, and stained for DNA using ProLong Diamond antifade mountant containing DAPI (50 μg/mL). Slides were then imaged using a Nikon Eclipse C2 laser scanning confocal system mounted on a Ti-E motorized inverted microscope. Excitation/stimulation was performed with a solid state 405/488/561/640 laser unit using a CFI Apo 60X oil immersion objective, NA 1.40. Images were collected with a high sensitivity C2-DU3 Detection Unit with 435/34, 525/50, and 600/50 filters. Confocal images were analyzed using Imaris x64 software version 10.1.0.

### Live cell imaging and time-lapse microscopy

A549 cells were seeded onto collagen-coated 35-mm glass-bottom dishes (MatTek) and cultured to confluence. Prior to infection, treatment, and imaging, cells were washed three times with PBS and incubated in infection medium supplemented with DAPI (50 μg/mL), wheat germ agglutinin–Alexa Fluor 555 (WGA-A555; 1 μg/mL), and CellEvent Caspase-3/7 Green Detection Reagent (2 μM; ThermoFisher). For apoptosis induction, cells were treated with staurosporine (1 or 10 μM; Sigma) for 7 h. For infection experiments, cells were exposed to Spn strains TIGR4 or EF3030 and incubated for 10 h at 37 °C in a humidified chamber (H301-Nikon-TI-S-ER) with 5% CO_2_ maintained by an Okolab controller. Live-cell confocal imaging and time-lapse acquisition were performed using a Nikon Eclipse C2 laser scanning confocal microscope with the specifications described above. Image processing and analysis of both static and time-lapse datasets were conducted using Imaris software (x64, version 10.1.0).

### Bulk RNA sequencing analysis

Human bronchial Calu-3 cells were seeded in 100 mm tissue culture treated dishes (Fisher) and incubate until they became polarized, 10-day post seeding. Cells were washed three times with sterile PBS and infected with Spn, mutant derivatives, or mock-infected, and incubated for 4 h at 37°C in a 5% CO_2_ atmosphere. After incubation, supernatant containing planktonic pneumococci were removed, the infected cells were harvested in one ml TRIzol (Invitrogen) and immediately flash frozen at -80°C. The following procedures were performed by the UMMC Molecular and Genomics Core Facility (www.umc.edu/genomicscore). RNA was extracted using the Pure Link RNA Mini Kit from archived samples (Invitrogen) according to manufacturer instructions and assessed for quality control parameters of minimum concentration and fidelity (i.e. 18S and 28S bands, RIS >8) through the UMMC Molecular and Genomics Core Facility (MGCF). Libraries were prepared for RNA sequencing per detailed protocols from Illumina mRNA stranded kit. Briefly, a pooled library was prepared from using n=12-24 samples per group using TruSeq mRNA Stranded Library Prep Kit (Set-A-indexes), quantified with the Qubit Fluorometer (Invitrogen), and assessed for quality and size Qiagen QIAxcel Advanced System or Agilent TapeStation. The library was sequenced using the NextSeq 2000 using P3 (200 cycles, paired end 100bp) on the Illumina NextSeq 2000 platform. Sequenced reads were assessed for quality using the Illumina BaseSpace Cloud Computing Platform and/or custom analysis pipeline and FASTQ sequence files were aligned reads to the reference genome (e.g., Ensembl/Hg38, Rnor_6.0, etc.) using DRAGEN RNA Pipeline Application (v.3.7.5) or custom analysis pipelines developed through MGCF. Differential expression was determined using DRAGEN Differential Expression Application (v.3.10.4) of DESeq2. Differentially expressed genes (DEGs) were identified using cutoffs of log2 (fold change) > 0.5 (∼1.4-fold) and log2 (fold change) > 1 (2- fold), with an adjusted *p*-value < 0.05. Gene Set Enrichment Analysis was performed using Reactome analysis tools (https://reactome.org/) or other databases. Data was deposited into NCBI GEO data under accession number GSE301601.

Gene expression study deposited in NCBI-GEO (GSE195778)(52) was utilized to investigate the expression of genes in the D39 background. We focused on RNA data from human bronchial epithelial cell line BEAS-2B were infected with Spn at multiplicity of infection (MOI) of 1 for 9 or 16 h, and compared against the mock-infected control.

To elucidate the biological implications of Spn infection, functional enrichment analysis was performed on differentially expressed genes (DEGs) using DAVID (v2023q1) and g:Profiler (v.e109_eg56_p17). DAVID and g:Profiler were used to identify enriched Gene Ontology (GO) terms across three categories: Biological Process (BP), Molecular Function (MF), and Cellular Component (CC). In TIGR4-infected Calu-3 cells, GO-BP analysis revealed significant enrichment of terms related to “mitochondrial organization” (GO:0007005, FDR = 2.3e-04), “cellular response to oxidative stress” (GO:0034599, FDR = 8.7e-03), and “apoptotic process” (GO:0006915, FDR = 1.2e-02), consistent with Spn-H_2_O_2_-induced mitochondrial dysfunction and limited apoptosis. GO-CC terms highlighted “mitochondrial inner membrane” (GO:0005743, FDR = 4.5e-03) and “respiratory chain complex” (GO:0098803, FDR = 9.1e-03), aligning with the observed inhibition of CIDR. KEGG pathway analysis via DAVID identified downregulation of the TCA cycle pathway (hsa00020, FDR = 3.8e-03) in TIGR4-infected cells, corroborating the inhibition of TCA cycle enzymes like ACO2 and OGDHC, while the “Glycolysis/Gluconeogenesis” pathway (hsa00010, FDR = 1.1e-02) was upregulated, reflecting the enhanced glycolysis observed (e.g., HK2 upregulation). Complementary analysis using g:Profiler provided additional insights into molecular networks. The “oxidative phosphorylation” pathway (KEGG:00190, p-adj = 2.4e-03) was significantly downregulated in TIGR4-infected cells, consistent with CIDR inhibition, while “response to reactive oxygen species” (GO:0000302, p-adj = 5.6e-03) was enriched, supporting the upregulation of oxidative stress response genes (SOD2, TXNRD2). Notably, in TIGR4Δ*spxB*Δ*lctO*-infected cells, these pathways showed minimal perturbation, underscoring the role of H_2_O_2_ in driving these changes.

### Heart mitochondria isolation

Intact rat heart mitochondria were isolated from Sprague Dawley rats similar to published methods (64, 65). Mitochondria were isolated from rat heart tissue due to their higher yield and robust respiratory performance compared to lung epithelial mitochondria, enabling reliable and high-quality bioenergetic analyses. While heart mitochondria preferentially utilize fatty acids and exhibit higher respiratory capacity, lung epithelial mitochondria rely more on glucose and glutamine and may have distinct ROS handling properties. These differences were considered in the context of modeling mitochondrial responses to Spn infection, with the conserved respiratory machinery supporting the relevance of our findings. In brief, heart tissue was minced with a razor blade in 500 µL of MSM buffer (220 mM mannitol, 70 mM sucrose, 5 mM MOPS pH 7.4) supplemented with 2 mg/mL bacterial proteinase type XXIV. The minced tissue was added to 5 mL of ice-cold isolation buffer (MSM buffer supplemented with 2 mM EDTA and 0.2% fatty acid-free BSA) and homogenized on ice with a glass homogenizer and a Teflon pestle for 4 strokes. To inhibit the proteinase, 0.1 mM phenylmethylsulfonyl fluoride (PMSF) was used. The tissue homogenate was centrifuged at 300×g for 10 min at 4°C. The supernatant was then centrifuged at 3000×g for 10 min at 4 °C, and the mitochondrial pellet washed once in ice-cold MSM buffer. Protein concentration was determined by using the BioRad DC protein assay.

### Hydrogen peroxide quantification

H_2_O_2_ levels were quantified from culture supernatants of different Spn strains. Each strain and culture condition was assessed in duplicate using 6-well plates and incubated for 4 hours at 37°C with 5% CO_2_. Cultures were then collected, centrifuged for 5 minutes at 21,130×g, and the supernatants were filter-sterilized using syringe filters (0.22 µm pore size). H_2_O_2_ was quantified using the Amplex Red H_2_O_2_ assay kit (Molecular Probes), according to the manufacturer’s instructions. Measurements were taken using an SYN H1 Hybrid Biotek reader (Agilent) or an Oroboros O2k FluoRespirometer(66).

### Purification of cytosolic and mitochondrial fractions and Western blotting

Calu-3 cells, infected as described, were incubated for 12 h at 37°C in 5% CO_2_, with mock- infected controls. At 12 h, 10 µM DCP-Rho1 (Cayman) was added for 10 min. Cells were washed twice with ice-cold PBS, scraped from 6-well plates, and lysed in RIPA buffer with 1x Complete Protease Inhibitor Cocktail EDTA-free (Roche) for 30 min on ice. Lysates were centrifuged (13,500 × g, 20 min, 4°C), and the supernatant (cytosolic fraction) was collected. Mitochondria were isolated from separate cells using the Mitochondria Isolation Kit for Cultured Cells (Thermo Fisher Scientific). Protein concentrations were measured with the BCA Protein Assay Kit (Thermo Fisher Scientific).

For Western blotting, equal protein amounts were mixed with 4× reducing sample buffer, boiled (5 min, 100°C), and separated on 12% Mini-PROTEAN TGX gels (Bio-Rad) at 90 V for 2 h. Proteins were transferred to nitrocellulose membranes using the Trans-Blot Turbo System (Bio-Rad) with 20% ethanol transfer buffer. Membranes were blocked (1 h, room temperature) in TBST (0.1% Tween 20) with 5% non-fat dry milk, then incubated overnight at 4°C with primary antibodies (anti-β-actin, Proteintech; anti-TRITC, Thermo Fisher Scientific; anti- MTCO2, Invitrogen) in TBST with 5% milk. After three 5-min TBST washes, membranes were incubated (1 h, room temperature) with StarBright Blue 520 anti-mouse or anti-rabbit secondary antibodies (1:2,500, Bio-Rad) in TBST with 5% milk, followed by six 5-min TBST washes. Blots were imaged on a ChemiDoc MP System (Bio-Rad) using Image Lab 5.0 software.

### Animal information

Sprague Dawley rats were purchased from Jackson Laboratory (Maine, US) at 7 weeks of age and allowed to acclimate for at least 1 week before use. All animals were housed in the Center for Comparative Research animal facilities of the University of Mississippi Medical Center (UMMC). Animals were kept on a 12 h light/12 h dark cycle and fed a standard laboratory rodent diet (Teklad 8640). All procedures were conducted in accordance with the National Institutes of Health’s Guide for the Care and Use of Laboratory Animals and approved by the UMMC Institutional Animal Care and Use Committee (Protocol #1582).

### Experiments of mitochondrial respiration and mitochondrial complex activity

Measurement of mitochondrial respiration follows previously published protocols(64, 65). In brief, Spn strains were cultured in THY for 4 h 37°C with a 5% CO_2_ atmosphere. Sterilized supernatants from each sample were then used to measure mitochondrial respiration driven through complex-I and II and activity through complexes I, II, III, and IV. In select experiments, catalase (to scavenge H_2_O_2_) or exogenous H_2_O_2_ (50 µM) was added to TIGR4 or TIGR4ΔspxBΔlctO supernatants to evaluate the reversibility of complex I inhibition. Complex-I driven respiration was measured through 20 mM glutamate plus 5 mM malate or 10 mM pyruvate plus 5 mM malate substrates with addition of 2 mM ADP. Complex-II driven respiration was measured through 20 mM succinate as the substrates with addition of 2 mM ADP. For fatty acid oxidation, the reaction contained 0.5 mM malate, 2 mM ATP, 1 mM NAD+, 25 µM cytochrome c, 0.1 mM coenzyme A, and either 0.5 mM oleate (long-chain fatty acid, LCFA) or 0.5 mM octanoate (medium-chain fatty acid, MCFA). All measurements were performed in an Oroboros O2K FluoRespirometer using a buffer containing 10 mM KPi, 5 mM MgCl2, 30 mM KCl, 1 mM EDTA, and 75 mM Tris, pH 7.5. All samples were normalized to protein content as determined by BioRad DC assay and reported as nmol e-/min/mg protein. *Complex I* activity was measured as the time-dependent oxidation of NADH at 340 nm using a measured extinction coefficient of 6.22 mM^-1^cm^-1^. The reaction mixture in a regular quartz cuvette contained 10 mM KPi, 5 mM MgCl_2_, 30 mM KCl, 1 mM EDTA, 75 mM Tris, pH 7.5, 100 µM Coenzyme Q1 (CoQ1), 5 µM antimycin A, and 5 µL heart mitochondria in a final volume of 2 mL. The reaction was initiated by adding 100 µM NADH and the fastest rates were obtained after NADH was added. *Complex II* activity was measured as the time-dependent oxidation of 2,6-dichloroindophenol sodium salt hydrate (DCPIP) at 600 nm using a measured extinction coefficient of 19.1 mM^-1^cm^-1^. The reaction mixture in a micro-cuvette contained 10 mM Kpi, 5 mM MgCl_2_, 30 mM KCl, 1 mM EDTA, 75 mM Tris, pH 7.5, 20 mM succinate, 80 µM DCPIP, 1 mM KCN, 5 µM Antimycin A, 4 µM rotenone, and 5 µL of heart mitochondria in a final volume of 200 µL. Reactions were initiated by adding 50 µM decylubiquinone and the linear rates were obtained. *Complex III* activity was measured as the time-dependent reduction of cytochrome *c* at 550 nm using a measured extinction coefficient of 18.7 mM^-1^cm^-1^. The reaction mixture in a regular glass cuvette contained 10 mM Kpi, 5 mM MgCl_2_, 30 mM KCl, 1 mM EDTA, 75 mM Tris, pH 7.5, 3 mM KCN, 4 µM rotenone, 60 µM cytochrome *c*, 5 µL of heart mitochondria in a final volume of 2 mL. *Complex IV* activity was measured as the time-dependent oxidation of N,N,N’,N’-Tetramethyl-p-phenylenediamine dihydrochloride (TMPD) to donate electrons to cytochrome *c*. The reaction mixture in a cuvette contained 50 mM Tris, 8 mM KCl, 1 mM EDTA, 16 μg/μL catalase, pH 7.4, 3 mM ascorbate, 0.3 mM TMPD, and 25 μM cytochrome c.

Reactions were initiated with 100 µM reduced decylubiquinone. The rate was linear for 1-2 min. *Aconitase* activity was measured as the time-dependent formation of cis-aconitate at 240 nm using a measured extinction coefficient of 2.2 mM^-1^cm^-1^ (67). The reaction mixture in a quartz cuvette containing 50 mM Tris–HCl (pH 7.4) containing 0.6 mM MnCl2, 30 mM citrate, and 0.2% peroxide-free Triton-X 100. For each enzyme, the inhibited non-enzymatic background rates were subtracted from the enzymatic rates. *Glutamate dehydrogenase* and α*- ketoglutarate dehydrogenase* were measured using the kits from Sigma (MAK099 and MAK189) following the provided protocols.

### Flow cytometry analysis using caspase-3/7 reagent and JC-1 assay

To assess mitochondrial membrane potential and apoptosis in human alveolar A549 cells infected with Spn, the MitoProbe JC-1 assay kit (Invitrogen) and the CellEvent Caspase-3/7 Green Detection Reagent (ThermoFisher) were employed. A549 cells grown in a 6-well plate were washed and treated with infection media as described above. Cells were then infected with Spn and incubated for 10 h. After incubation, cells were lifted from the wells using 0.25% Trypsin-EDTA, counted, and ∼1×10^6^ cells were used for JC-1 staining following the manufacturer’s protocol. For caspase-3/7 detection, cells were incubated with the CellEvent Caspase-3/7 Green Detection Reagent (2 µM final concentration) for 30 minutes at 37°C prior to analysis, following the manufacturer’s instructions. As a control, uninfected cells were treated with 125 μM Carbonyl cyanide m-chlorophenyl hydrazone (CCCP) for 10 minutes or 2 mM H_2_O_2_ for 1 h to induce mitochondrial depolarization or oxidative stress, respectively. In addition, staurosporine (Sigma, 1 µM and 10 µM, 4 hours) was used as a positive control for caspase-3/7 activation. Subsequently, cells were stained with 1 μM SYTOX Blue Dead Cell Stain (Invitrogen) for dead cells. Flow cytometry was performed on a NovoCyte (Agilent). JC-1 was excited with a 488 nm laser, and the emission was collected in the PE-Texas Red (red, 615/20) and FITC (green, 530/30) channels, to assess mitochondrial membrane potential.

Caspase-3/7 activation was detected in the FITC channel (530/30 nm filter) following excitation at 488 nm. SYTOX Blue Dead Cell Stain was excited by the violet 405 nm laser and detected in the 530/30 nm channel. Fluorescence compensation, including correction for JC-1 spectral overlap between FITC and Texas Red channels, was performed using FMO and single-color controls. Data from 50,000 events per sample were acquired and analyzed using NovoExpress software (Agilent).

### TUNEL assay

To identify late-stage apoptosis in Spn-infected human alveolar A549 cells, the Click-iT Plus TUNEL assay kit (Invitrogen) was utilized. A549 cells were seeded in a 6-well plate with glass coverslips and used after ∼7-10 days post-seeding. Before infection, A549 cells were washed and treated with infection media as described above. Cells were then infected with the indicated Spn strain and incubated for 8 h. As a control, some cells were treated with 2 mM H_2_O_2_ every 2 h for 8 h. After incubation, cells were stained following the manufacturer’s protocol. Following staining, cells were washed, coverslips were removed, and the cells were allowed to dry before being mounted and stained for DNA using ProLong Gold Antifade Mountant containing DAPI. Slides were then imaged using a Nikon A1R HD25 laser scanning confocal upright microscope. Confocal images were analyzed using Imaris x64 software version 10.1.0.

### Activity of glutamate dehydrogenase in bronchial Calu-3 cells infected with pneumococcal strains

Calu-3 cells grown in 100 mm tissue culture-treated dishes (Fisher) were washed three times with sterile PBS to remove antibiotics and then added with 10 ml of infection medium, consisting of EMEM supplemented with 1% HEPES buffer and 5% fetal bovine serum (FBS). Cells were either mock-infected or infected with TIGR4 or TIGR4Δ*spxB*Δ*lctO* and incubated for 8 h at 37°C in a 5% CO_2_ atmosphere. The supernatant, containing the planktonic bacterial fraction, was collected for HPLC studies, as detailed below. Infected cells were washed once with PBS and harvested using a cell scraper in 1 ml of cold sterile PBS. The cells were centrifuged at 300 × g for 5 minutes, and the resulting pellet was resuspended in 200 μl of PBS. The cells were then lysed using a Dounce homogenizer, and the lysate volume was adjusted to 600 μL with sterile PBS. The lysate was centrifuged at 10,000 × g for 15 minutes at 4°C, and the soluble fraction was used to assess the enzymatic activity of glutamate dehydrogenase (GDH) and for HPLC analysis. Assays were performed in technical triplicates with two biological replicates.

### Glutamate dehydrogenase activity and metabolite quantification in Spn-infected human respiratory cells

Human bronchial Calu-3 cells were cultured for 10 days in 55 cm² Petri dishes in Eagle’s Minimum Essential Medium (EMEM) supplemented with 1% HEPES buffer and 5% fetal bovine serum (FBS). Cells were washed three times with sterile phosphate- buffered saline (PBS) to remove antibiotics and overlaid with 10 ml of infection medium (EMEM with 1% HEPES and 5% FBS). Cells were infected with Spn strains and incubated at 37°C for 8 h. Mock-infected cells served as controls. For secreted metabolite analysis, 1 ml of infection medium was centrifuged at 10,000 × g for 5 min to remove planktonic bacteria, and the supernatant was collected. Cells were washed once with PBS, scraped in 1 ml PBS, and centrifuged at 300 × g for 5 min. For glutamate dehydrogenase (GDH) activity, the cell pellet was resuspended in 200 µl PBS, homogenized using a Dounce homogenizer, diluted with 400 µl PBS, and centrifuged at 10,000 × g for 15 min. The resulting supernatant (cell extract) was used for GDH activity assays and intracellular metabolite quantification. GDH activity was measured using a non-radioactive colorimetric assay (MAK499-1KT, Sigma-Aldrich, St. Louis, MO) according to the manufacturer’s instructions. For metabolite quantification, lactate, acetate, and citrate levels in cell-free supernatants (extracellular) and cell extracts (intracellular) were analyzed by high-performance liquid chromatography (HPLC) using a Thermo Dionex Ultimate 3000 UHPLC System equipped with an Aminex HPX-87H column (Bio-Rad, Hercules, CA) maintained at 25°C. An isocratic elution was performed with 5 mM H_2_SO_₄_ at a flow rate of 0.6 mL/min, and metabolites were detected at 208 nm. Samples were prepared by mixing 100 µl of supernatant or cell extract with 100 µl of 95% ethanol, incubating at –80°C for 15 min to precipitate proteins, and centrifuging at 20,000 × g for 15 min. A 20 µl aliquot of the resulting supernatant was injected for analysis. All assays were performed with two technical triplicates and three biological replicates. Statistical analyses are described in the relevant figure legends.

### Statistical analysis

We performed one-way analysis of variance (ANOVA) followed by Dunnett’s multiple-comparison test when two or more groups were involved. All analysis was performed using GraphPad Prism software (version 10.2.1).

## ACKNOWLEDGEMENTS

This study was supported in part by grants from the National Institutes of Health (NIH), including R21AI144571, R01AI175461 (to J.E.V.) and R01GM142113 and RR17767 (to K.W.).

Confocal microscopy and mitochondrial respiration studies (K.E.) were supported by the National Institute of General Medical Sciences (NIGMS) under Award Number P20GM121334.

J.E.V. is also supported by NIGMS through the Molecular Center of Health and Disease (P20GM144041). RNA-seq data analysis was performed at the UMMC Molecular and Genomics Facility, supported by funds from NIGMS (P20GM144041), Mississippi INBRE (P20GM103476), the Obesity, Cardiorenal, and Metabolic Diseases COBRE (P30GM149404), and the Mississippi Center of Perinatal Research (P20GM121334). The content is solely the responsibility of the authors and does not necessarily represent the official views of the NIH or the U.S. Department of State. The authors thank Dr. David Brown (UMMC) for his assistance with confocal microscopy.

